# A gradient border model for cell fate decisions at the neural plate border

**DOI:** 10.1101/2022.02.15.480567

**Authors:** Alexandre Thiery, Ailin Leticia Buzzi, Eva Hamrud, Chris Cheshire, Nicholas Luscombe, James Briscoe, Andrea Streit

## Abstract

The vertebrate ‘neural plate border’ is a transient territory located at the edge of the neural plate containing precursors for all ectodermal derivatives: the neural plate; neural crest; placodes; and epidermis. Elegant functional experiments in a range of vertebrate models have provided an in-depth understanding of gene regulatory interactions within the ectoderm. However, these experiments conducted at tissue level raise seemingly contradictory models for fate allocation of individual cells. Here, we carry out single cell RNA sequencing of chick ectoderm from primitive streak to neurulation stage, to explore cell state diversity and heterogeneity. We characterise the dynamics of gene modules containing key factors known to regulate ectodermal cell fates, allowing us to model the order in which these fates are specified. Furthermore, we find that genes previously classified as neural plate border specifiers typically exhibit dynamic expression patterns and are biased towards either placodal or neural crest fates, revealing that the neural plate border should be seen as an anatomical region of the ectoderm and not a discrete transcriptional state. Through co-expression of placodal and neural crest markers, we identify a population of border located unstable progenitors (BLUPs) which gradually reduces in size as fate segregation occurs. Considering our findings, we propose a ‘gradient border’ model for cell fate choice at the neural plate border, with the probability of cell fate allocation closely tied to the spatiotemporal positioning of cells.

## Introduction

The vertebrate nervous system is arguably the most complex organ of our body. During development, it arises from only three progenitor populations: the neural plate generates the entire central nervous system, while neural crest and sensory placodes jointly form the peripheral nervous system. Placode derivatives are confined to the head contributing to sense organs and cranial ganglia, while neural crest cells form along the entire body axis to generate various cell types including neurons and glial as well as parts of the craniofacial skeleton, cartilage and teeth (for review: Baker & Bronner-Fraser, 1997a; Baker & Bronner-Fraser, 2001; Grocott et al., 2012; Patthey et al., 2014; Pla & Monsoro-Burq, 2018; Prasad et al., 2019; Simoes-Costa & Bronner, 2015).

The neural plate becomes morphologically distinct shortly after gastrulation, although neural, neural crest and placode precursors continue to be intermingled at its edge, in a territory termed the neural plate border (NPB). The NPB has been characterised by the co-expression of neural and non-neural markers as well as so-called NPB specifiers (Pla & Monsoro-Burq, 2018; Roellig et al., 2017; Streit, 2002; Thiery et al., 2020). As the neural plate invaginates, placode precursors remain in the surface ectoderm, while neural crest cells are largely incorporated into neural folds from where they delaminate and migrate extensively. Although the induction of these three fates has been studied for decades, some of the most fundamental questions remain unresolved. For example, we do not know when and how these lineages segregate, whether they do so in a specific order and how individual cells make these decisions.

Modulation of different signalling pathways plays a crucial role in controlling the fate of all ectodermal cells. FGF signalling is required for the specification of neural, neural crest and placode precursors and for positioning the NPB (Ahrens & Schlosser, 2005; Litsiou et al., 2005; Londin et al., 2005; Monsoro-Burq et al., 2003; Streit et al., 2000; Streit & Stern, 1999; Stuhlmiller & García-Castro, 2012; Wilson et al., 2000; Yardley & Garcia-Castro, 2012). While BMP signalling must be inhibited for neural development (Hemmati-Brivanlou et al., 1994; Linker & Stern, 2004; Londin et al., 2005; Sasai et al., 1995), it is necessary for placode precursor formation at gastrulation, but must be switched off thereafter (Ahrens & Schlosser, 2005; Kwon et al., 2010; Litsiou et al., 2005). Likewise, neural crest cells need BMP activity early, but unlike placodes continue to do so as they are specified (Barth et al., 1999; Kwon et al., 2010; Liem et al., 1995; Marchant et al., 1998; Steventon et al., 2009). Finally, Wnt activity is required for neural crest cell induction (Garcia-Castro et al., 2002; Steventon et al., 2009) but must be inhibited for neural and placode precursors to form (Heeg-Truesdell & LaBonne, 2006; Litsiou et al., 2005; Patthey et al., 2008). These signals activate different transcription factor networks in a temporally and spatially controlled manner, which in turn are thought to impart cell identity and have therefore been termed fate ‘specifiers’. For example, *Msx1* and *Pax3* have been implicated as NPB specifiers (Milet et al., 2013; Monsoro-Burq, 2015; Plouhinec et al., 2014), while *Pax7* (Basch et al., 2006) and *Six1* (Brugmann et al., 2004; Chen et al., 2009; Christophorou et al., 2009; Ozaki et al., 2004) are involved in neural crest and placode precursor specification, respectively. These factors appear to act in feed forward loops, upregulating their own expression while simultaneously repressing factors that specify alternative fates (for review: Grocott et al., 2012; Thiery et al., 2020).

Based on these studies combined with tissue transplantation experiments, two apparently contradictory models for the segregation of neural, neural crest and placodal fates have emerged (for review: Schlosser, 2006, 2014; Thiery et al., 2020). The ‘binary competence model’ suggests that an initial step in lineage segregation entails a subdivision of the ectoderm into neural/neural crest and placode/epidermal territories (Ahrens & Schlosser, 2005; Pieper et al., 2012; Schlosser, 2014). This implies a closer relationship of neural crest cells with the central nervous system and of placodes with the epidermis. In contrast, the ‘NPB model’ suggests that cells at the border of the neural plate have mixed identity and retain the ability to generate all ectodermal derivatives until after neurulation begins (Baker & Bronner-Fraser, 1997b; Roellig et al., 2017; Streit & Stern, 1999). Studies in *Xenopus* have taken this idea a step further, revealing that unlike other ectodermal cells, cells at the NPB retain pluripotency markers which is key for their multipotency (Buitrago-Delgado et al., 2015; Prasad et al., 2012). In support of this idea, the NPB region is initially characterised by co-expression of neural and non-neural markers (for review: Grocott et al., 2012; Pla & Monsoro-Burq, 2018; Thiery et al., 2020). Fate specifier genes are activated within this territory in partially overlapping patterns and resolve into distinct domains as neurulation begins. While both models are attractive and provide working hypotheses, the underlying experiments were analysed at cell population, rather than at single cell level. They therefore do not address how individual cells at the NBP make decisions, nor whether they are multipotent or a population of mixed progenitors that are already biased towards their later identity. Recent findings have proposed that prior to fate commitment, cells show increasing levels of heterogeneity in gene expression and do not exhibit a definitive transcriptional signature (Soldatov et al., 2019). It is therefore possible that cells at the NPB include such undetermined progenitor cells.

Studies in chick have begun to characterise cell heterogeneity at the NPB at a single cell level revealing that factors previously thought to be fate-specific are co-expressed in individual cells even at late neural tube stages (Roellig et al., 2017; Williams et al., 2022). Furthermore, lineage tracing using a ‘neural-specific’ Sox2 enhancer shows that Sox2^+^ cells can contribute not only to neural tissue but also to neural crest and epidermis (Roellig et al., 2017). However, these studies only surveyed a relatively small number of ectodermal cells and, focusing on neural crest cells, did not include data for definitive placodal cells. It is therefore challenging to assess how these fates segregate. Nevertheless, these findings suggest that cell fate allocation at the NPB occurs over a much longer developmental period than previously thought and that NPB cells retain plasticity until early neurulation. However, the underlying molecular mechanisms of how individual cells make decisions at the NBP remain poorly understood.

Here, we explored the transcriptional changes as cells transit from the neural plate border at early primitive streak to definitive neural, neural crest and placodal fates at single cell level (for analysis workflow see Figure 1). In total we present single cell RNAseq (scRNAseq) data for 17,992 high quality cells. Rather than using 2-3 molecular markers to assign cell identity, we established a binary knowledge matrix using previously published *in situ* hybridisation expression data. This allows us to classify cells at each developmental stage, despite the high levels of heterogeneity observed. We find that prior to head fold stages cells do not fall into transcriptional distinct groups but are characterised by graded and partially overlapping gene expression. An early cell population located at the NPB (eNPB) can be identified by co-expression of neural and non-neural genes prior to the upregulation of fate specifiers. As development proceeds, ectodermal cell diversity increases, with a division of neural and non-neural fates becoming apparent at the first somite stage together with the emergence of neural crest cells. By the 8-somite stage, neural, neural crest and placodal cells are largely segregated, although even at this late stage we find cells with heterogenous expression profiles.

Using a gene module approach, we identify dynamic changes in groups of genes that are characteristic for neural, neural crest and placodal fates. Comparing co-expression of whole gene modules (rather than individual transcripts) reveals that many cells at the NPB co-express gene modules which later become restricted to placodal or neural crest lineages. These cells may have the potential to give rise to both (or more) fates and we therefore refer to them as ‘Border-Located Unstable Progenitors’ (BLUPs). Interestingly, as progenitors segregate into their respective fates, we no longer identify NPB clusters but continue to find BLUPs. BLUPs connect segregating cell populations, resembling previously characterised bridge cells in the differentiating murine neural crest which exhibit high heterogeneity prior to segregation (Soldatov et al., 2019). Here, we propose that BLUPs continue to be plastic until neural tube closure, maintaining the ability to contribute to different fates. Overall, our analysis highlights that although the NPB demarcates an anatomical region, it is not a discrete transcriptional state. Instead, we identify BLUPs which we predict to be in an undetermined state and to retain the potential to contribute towards alternative ectodermal fates.

## Results

### Classification of ectodermal cells using a binary knowledge matrix of known marker genes

Neural, neural crest and placode cells arise from the embryonic ectoderm. However, how individual progenitors acquire their terminal fates and the sequence and timing of their segregation remain to be elucidated. To capture cells during this process, we carried out 10x Genomics single-cell mRNA sequencing (scRNAseq) on cells taken from a broad region of the chick epiblast at six developmental stages: HH4^-^/4 (primitive streak), HH5 (head process), HH6 (head fold), HH7 (1 somite), somite stage (ss) 4 and ss8 (Figure 2A). These stages were chosen because they encompass the initial formation of the neural plate to the beginning of neurulation when these fate decisions are thought to take place. After integration across batches, stringent quality control and removal of contaminating cell populations (see methods and Figure 2–figure supplement 1A-I), we captured a total of 17,992 high quality cells with approximately 2500-4000 cells per stage and an average of 3924 genes per cell. Dimensionality reduction and UMAP embedding reveals that cells at later stages displayed higher cell diversity compared with earlier stages where most cells were transcriptionally similar (Figure 2B-E and Figure 3A-D).

**Figure 2.**
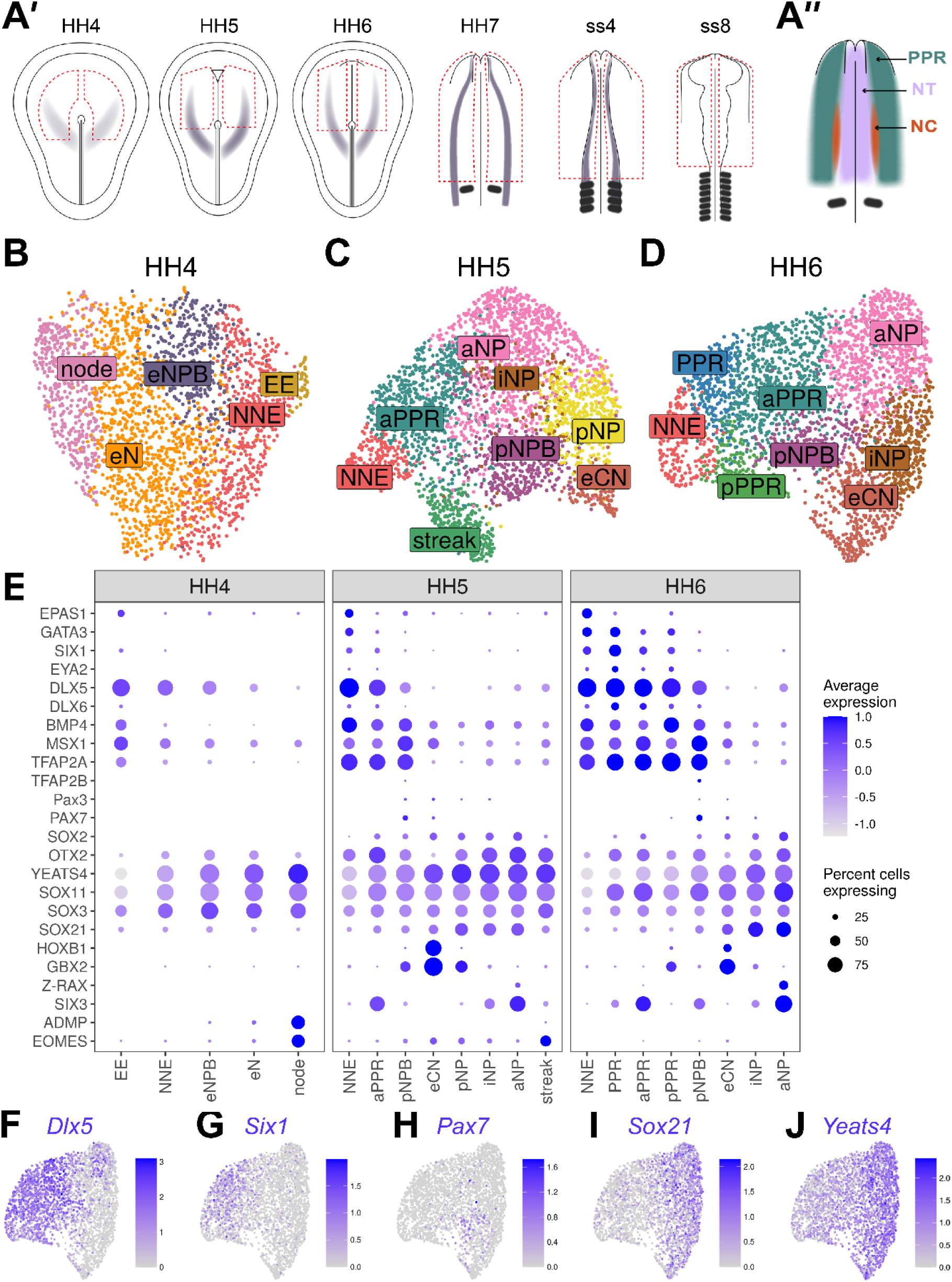
Cells at stages HH4 to HH6 reflect the anterior-posterior and medio-lateral axes in the embryo. **(A’)** Dorsal view schematics of chick embryo at stages HH4-ss8 depicting the ectodermal tissue region dissected for 10x scRNAseq (red dotted line). Purple shading illustrates previously characterized region of *Pax7* expression (Basch et al., 2006). **(A’’)** Schematic of a 1-somite stage (HH7) chick embryo illustrating the pre-placodal region (PPR), neural crest (neural crest) and neural tube (NT). **(B-D)** UMAP plots for cells collected at HH4^-^/4 (primitive streak), HH5 (head process), HH6 (head fold) stages. Cell clusters are coloured and labelled based on their semi-unbiased cell state classifications (binary knowledge matrix available in Supplementary File 1). **(E)** Dot plots displaying the average expression of key marker genes across cell states at HH4, HH5 and HH6. The size of the dots represents the number of cells expressing a given gene within each cell population. **(F)** Feature plots revealing the mediolateral expression of key marker genes at HH6 and their overlapping expression.

**Figure 3.**
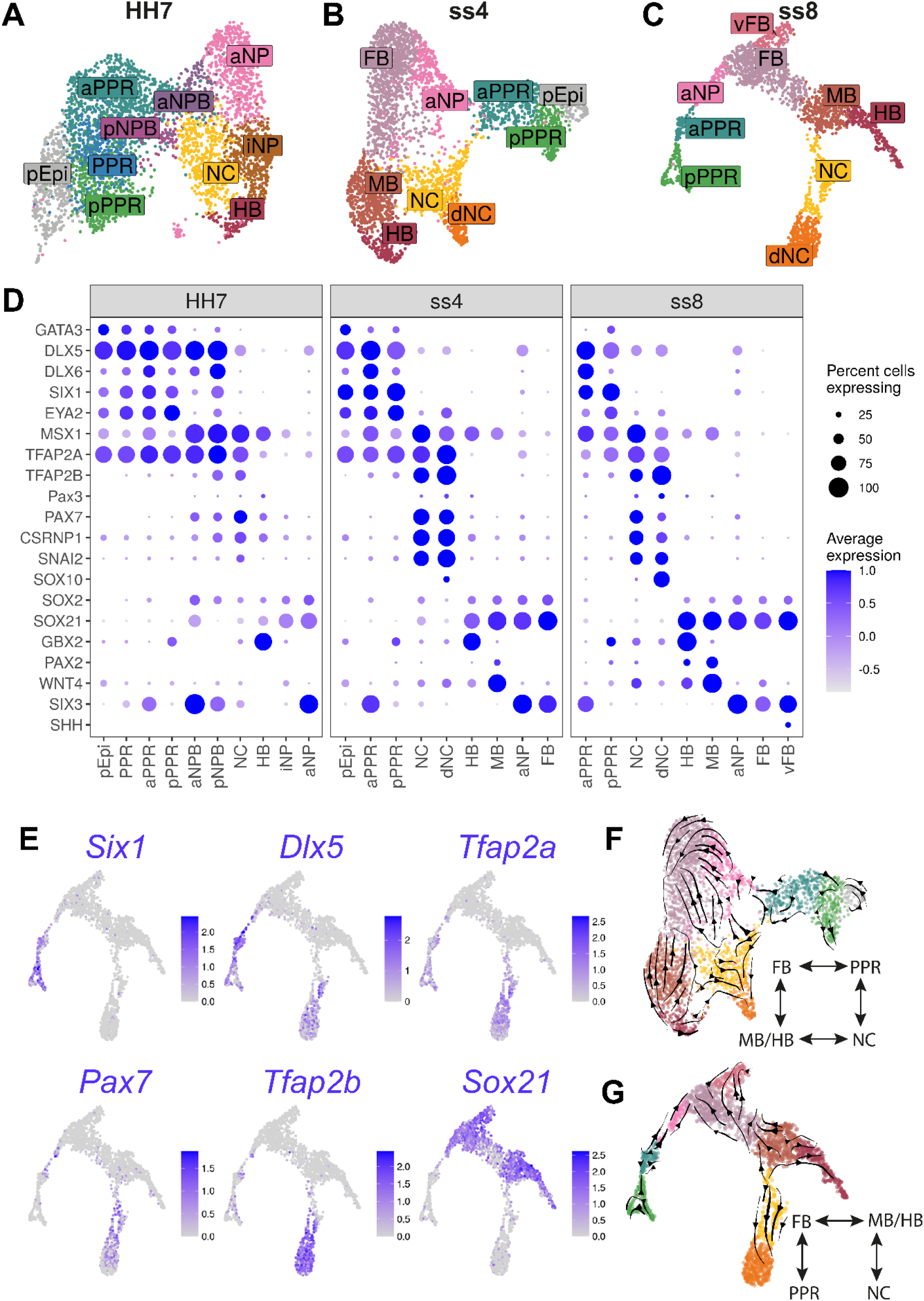
Increased cell diversity and lineage segregation from HH7 to ss8. **(A-C)** UMAP plots for cells collected at HH7 (1 somite stage), ss4 (4 somite stage) and ss8 (8 somite stage). Cell clusters are coloured and labelled based on their semi-unbiased cell state classifications (binary knowledge matrix available in Supplementary File 1). **(D)** Dot plots displaying the average expression of key marker genes across cell states at HH7, ss4 and ss8. The size of the dots represents the number of cells expressing a given gene within each cell population. **(E)** Feature plots depicting the lineage restricted expression of neural (*Sox21*), placodal (*Six1, Dlx5, Tfap2a*) and neural crest (*Pax7, Tfap2a, Tfap2b*) markers. **(F-G)** UMAP plots for ss4 and ss8 overlaid with RNA velocity vectors depicting the predicted directionality of transcriptional change. Cells are coloured by cell state classification (shown in B and C). Schematics summarise the key fate segregation events taking place at these stages predicted by RNA velocity analysis.

Typically, scRNAseq cell states are classified using a small number of definitive marker genes which exhibit mutually exclusive expression patterns. However, this approach is difficult to apply to the NPB for several reasons. Firstly, the early NPB cannot be classified using definitive markers as it emerges in the region of overlap between neural and non-neural genes. Secondly, cells at the NPB are thought to be in a transitory state before they acquire their definitive ectodermal fate (neural, neural crest, placodal and epidermal). Finally, cells at the NPB appear to be highly heterogeneous, with precursors for the different fates found intermingled, even at late stages of neurulation (Bhattacharyya et al., 2004; Roellig et al., 2017; Streit, 2002; Williams et al., 2022). To overcome these challenges, we categorised cells using a binary knowledge matrix based on expression data obtained from the literature (Supplementary File 1). We examined available *in situ* hybridisation data of 76 genes with known regionalised expression patterns and binarized their expression within each defined cell state. This allowed us to define a total of 24 cell states spanning primitive streak to neural tube stages. Using this approach, we classify NPB cells not only by the expression of NPB specifiers *Pax7* and *Tfap2a*, but also by the overlapping expression of early neural and non-neural genes.

We first unbiasedly clustered cells from each developmental stage at a high resolution to ensure capturing the full diversity of cell states (Figure 2–figure supplement 1J). Clusters were then classified based on the similarity between their gene expression and the knowledge matrix (see methods). This classification approach successfully identified 22 distinct cell states within the ectoderm across all developmental stages. Early NPB and early neural cell populations can already be identified at primitive streak stages, while placode progenitor and neural crest states emerge at head process and ss4, respectively (Figure 2B and Figure 3A-B).

### Cell states emerging from primitive streak to head fold stages

To explore the temporal changes in cell diversity within the embryonic ectoderm, we compared cell states present at different stages and identified the time point when each emerged. We first examined stages HH4-HH6 to characterise events as the neural plate is established and placodal and neural crest cells are thought to be specified.

#### An early NPB population is identified at primitive streak stages prior to the expression of fate specifiers

By primitive streak stage, mediolateral (M-L) gradients of molecular markers demarcate pre-neural and non-neural territories within the chick epiblast (Pera et al., 1999; Rex et al., 1997; Sheng & Stern, 1999; Streit et al., 2000; Streit et al., 1998). At HH4, unbiased clustering clearly organises cells in a pattern reminiscent of this M-L axis (Figure 2B; from left to right) as evidenced by the expression of node (*ADMP*), pre-neural (*Yeats4, Sox3, Sox11*) and non-neural (*Dlx5/6, Bmp4, Msx1, Tfap2a*) genes (Figure 2E and Figure 2–figure supplement 2A). Cell states can be broadly characterised based on combinatorial gene expression (Figure 2E) and we identify five cell states: node, early neural (eN), early neural plate border (eNPB), non-neural ectoderm (NNE) and extraembryonic ectoderm (EE) (Figure 2B). While some genes are shared between many cells, early neural plate cells are defined by high levels of pre-neural transcripts while the non-neural and extraembryonic ectoderm are characterised by increased non-neural gene expression. Notably, we identify that an early NPB population which co-expresses both neural and non-neural genes. Thus, early NPB cells are already apparent prior to the differential upregulation of genes previously characterised as NPB specifiers (*Msx1, Pax7, Tfap2a, Dlx5/6, Pax3*) (Basch et al., 2006; Ezin et al., 2009; Hong & Saint-Jeannet, 2007; McLarren et al., 2003; Rothstein & Simoes-Costa, 2020; Streit & Stern, 1999) (for review: Simoes-Costa & Bronner, 2015; Thiery et al., 2020).

#### The first emerging NPB and PPR cells have regional identity

At HH5, cell positions in the UMAP continue to reflect organisation along the M-L axis in the embryo (Figure 2C), but new transcriptional states emerge. Whilst non-neural ectoderm cells display a similar expression profile at HH5 compared to HH4, neural plate cells differ from early neural cells in their expression of definitive neural (e.g., *Sox2*) and regional markers (Figure 2E; Figure 2–figure supplement 2B). Like at HH4, many cells continue to co-express neural and non-neural transcripts (Figure 2–figure supplement 2B). Within the latter cell group, we observe the upregulation of PPR genes (*Six1, Eya2*; Figure 2C, E: aPPR, anterior PPR) and for the first time NPB specifiers (*Msx1, Pax3, Pax7*; Figure 2C, E: pNPB, posterior NPB). These observations correlate well with gene expression in the embryo. *Msx1* is initially expressed in the extraembryonic and non-neural ectoderm, and together with *Pax7* begins to be upregulated at the NPB thereafter (Basch et al., 2006; Streit & Stern, 1999), while PPR markers are first expressed at HH5. Notably, the NPB initially has a posterior character, while placode precursors have anterior identity based on their expression of *Gbx2* and *Six3*, respectively (Figure 2E; see also below).

#### Anteroposterior organisation of cells at head process and head fold stages

Cells with distinct anteroposterior (A-P) character are first observed at HH5 through the restricted expression of various homeobox transcription factors. *Rax* and *HoxB1* are highly confined to distinct cell groups, whilst in contrast *Six3* and *Gbx2* exhibit broader negatively correlated expression gradients across cell clusters with some degree of overlap (Figure 2E and Figure 2–figure supplement 2B). The organisation of cells within the UMAP reflects the A-P expression gradients of these genes in the embryo (Chapman et al., 2002; Hidalgo-Sanchez et al., 2005; Ohuchi et al., 1999; Paxton et al., 2010). The overlap of A-P factors was used to distinguish different NP, PPR and NPB regions during unbiased cell state classification (Supplementary File 1).

Within the NP we can distinguish intermediate neural plate (iNP) states from the anterior and posterior neural plate (aNP, pNP) and from early caudal neural (eCN) states. However, within the PPR and NPB we identify only anterior (aPPR; *Six3^+^*) and posterior (pNPB; *Gbx2+*) cell states, respectively. While in UMAPs, aPPR cells are located between the NNE and anterior NP, the pNBP cluster resides between the iNP and pPPR clusters (Figure 2C, E Figure 2–figure supplement 2C), reflective of both its A-P and M-L position *in vivo*.

Between HH5 and HH6, posterior *Gbx2*^+^ PPR (pPPR) cell states emerge (Figure 2D), as do PPR cells that appear to lack regional identity. At this stage, *Otx2* and *Gbx2* expression domains in the embryo abut (Steventon et al., 2012), suggesting that cells with generic PPR character might be intermingled with those already biased towards axial identity and either retain the potential to contribute to multiple axial levels or will later become intermediate PPR cells.

Together, our data show that the emergence of cells with divergent transcriptional profiles resembles patterning of the ectoderm *in vivo*. However, unbiased cell clustering forces cells into discrete clusters which may not fully reflect heterogeneity in gene expression and cell diversity. Indeed, our single cell data reveal the heterogenous nature of gene expression: markers are broadly expressed and are not confined by cluster boundaries or by neighbouring expression domains (Figure 2F-J, Figure 2–figure supplement 2C). Despite this heterogeneity, our cell classification approach successfully identifies increased ectodermal cell diversity over time, with neural, posterior NPB and anterior PPR cells emerging when the definitive neural plate forms. Importantly, the relative positions of these cell states in the UMAPs reflect their location in the embryo.

### Increasing cell diversity from 1-somite to 8-somite stages

At HH7, the neural folds begin to elevate indicating the start of neurulation, while during early somitogenesis neural crest cells start to migrate and placode precursors begin to diversify. We therefore sought to characterise the transcriptional changes at single cell level as neural, neural crest and placodal fates emerge.

Following clustering, we identified two major superclusters at HH7, one containing neural tube and neural crest and the other placodal and future epidermal clusters (Figure 3A). These superclusters appear to be largely separate, both by their UMAP embedding and the restricted expression of definitive fate markers (Figure 3A, D). Whereas at earlier stages many cells co-express non-neural and neural markers (Figure 2E, F, J), at HH7 cells expressing non-neural/placodal (*Dlx5/6, Six1*) and neural transcripts (*Sox21, Sox2*) are largely confined to their respective supercluster (Figure 3–figure supplement 1A). It is at this stage that anterior NPB cells can first be identified transcriptionally, and together with posterior NPB cells they connect the two superclusters (Figure 3A; aNPB, pNPB).

#### The neural crest begins to be specified at early neurulation

Adjacent to the NPB clusters, we observe a neural crest cell cluster expressing early definitive neural crest markers including *Tfap2b, Snai2* and *Axud1 (Csrnp1*) alongside NPB markers (*Msx1*, *Pax7* and *Tfap2a*) (Figure 3D; NC and Figure 3–figure supplement 1A). Thus, neural crest cells first emerge as a distinct cell population at HH7. Although many genes are shared between the NPB and neural crest cell clusters, NPB cells also express PPR and non-neural markers (*Six1, Eya2* and *Dlx5/6*) which are in turn downregulated in neural crest cells (Figure 3D and Figure 3–figure supplement 1A). Furthermore, we find that *Msx1* and *Pax7* are not strongly expressed within placodal cell clusters. It is therefore possible that these markers do not represent NPB specifiers, but rather an early bias towards neural crest fate.

#### Neural, placodal and neural crest lineages are largely segregated by ss4

At ss4 and ss8, a NPB cell cluster can no longer be identified using the above criteria. Instead, we observe an increasing diversity of cells classified as placodal, neural crest or neural (Figure 3B, C, E). Within the neural cell population, forebrain, midbrain, and hindbrain (FB, MB, HB) clusters are clearly distinguishable. *Six3* and *Gbx2* expression define cells with fore- and hindbrain identity, respectively, whilst the upregulation of the dorsal midbrain marker *Wnt4 (Hollyday et al., 1995*) and mid/hindbrain boundary marker *Pax2* (Hidalgo-Sanchez et al., 1999) highlights midbrain-like cells (Figure 3D and Figure 3–figure supplement 1B, C). As development progresses, neural, neural crest and placode cells become increasingly distinct, with neural crest cells beginning to express a suite of definitive neural crest markers, including *Sox10* in the delaminating neural crest (dNC) cluster (Figure 3D).

Despite the apparent segregation of neural, neural crest and placodal fates at ss4 and ss8, small cell populations connect PPR, neural crest, forebrain and midbrain clusters, in a four-way pattern reflecting their A-P and M-L segregation *in vivo* (Figure 3B, C). To predict the directionality of transcriptional change within these populations, and hence their potential developmental trajectory, we performed RNA velocity analysis (Bergen et al., 2020; La Manno et al., 2018). This analysis predicts that they move away from cluster boundaries, towards definitive cell states (Figure 3F, G).

Together our data show that whereas cell states at HH4-HH6 exhibit broad overlapping gene expression, definitive fate markers are upregulated from HH5 onwards and subsequently gene expression is refined at the boundaries between cell clusters. At HH7 and ss4, placodal, neural crest and neural fates begin to segregate. However, even as the neural tube closes, cells continue to straddle cluster boundaries; these cells may be transcriptionally unstable and therefore still actively undergoing fate decision processes.

### Dynamic changes in gene module expression over time

Many transcriptional regulators and their interactions have been implicated in cell fate decisions as the ectoderm is specified and differentiates into its derivatives. However, studies have largely taken a gene candidate approach. To explore the full transcriptional dynamics of both known and new transcriptional regulators during the specification of neural, neural crest and placodal cells, we modelled the dynamics of whole gene modules rather than individual candidate genes. This allows us to investigate transcriptional dynamics in an unbiased manner whilst simultaneously minimising the loss of potentially important factors.

#### Defining neural, neural crest and placodal gene modules

To investigate the segregation of neural, neural crest and placode fates, we grouped together cells from all developmental stages (Figure 4A, B) and sought to identify gene modules that characterise each cell lineage. Gene modules are groups of genes that display similar expression profiles across a dataset. We calculated gene modules using the Antler R package (Delile et al., 2019), which iteratively clusters genes hierarchically based on their gene-gene Spearman correlation. This approach reveals 40 modules each showing highly correlated gene expression across all stages (HH4-ss8). To focus on gene modules which may include candidates for fate segregation, we included only those that are differentially expressed between any cell state and the rest of the dataset, and then filtered to remove those with differential expression between sequencing batches. Finally, since our data show that the segregation of neural, neural crest and placodal cell states is almost complete at ss8 (Figure 3C), we kept those gene modules that show differential expression between at least one of the three fates at ss8 (see methods). This resulted in nine modules that were considered further (Figure 4–figure supplement 1A and Figure 4–Source Data 1). Of these, four gene modules (GM21-24) are specifically expressed in the PPR at ss8, one (GM13) in neural crest cells, and the remaining four (GM7, 9, 11, 20) are upregulated primarily in neural clusters (Figure 4–figure supplement 1A).

**Figure 4.**
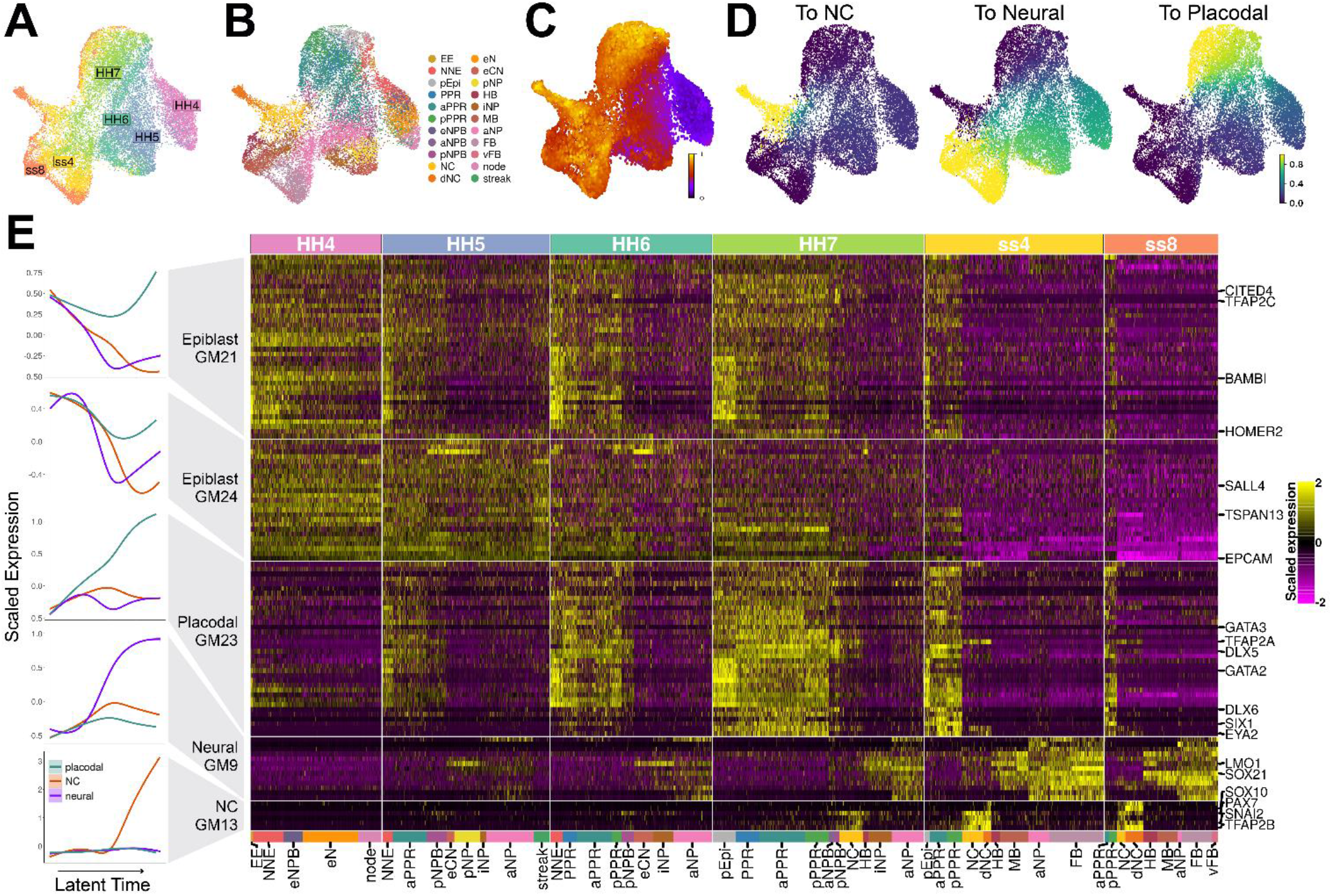
Gene module dynamics reveal key differences between the segregation of the PPR, neural crest and neural fates. **(A)** UMAP plot for the full dataset (HH4-ss8 combined) coloured and labelled by stage. **(B)** UMAP plot for the full dataset coloured by cell state classifications. Cell state classifications were calculated independently for each stage and transferred across for visualisation of the full dataset. **(C)** UMAP plot of the full dataset (HH4-ss8) showing cell latent time values. **(D)** UMAP plots of the full dataset showing the fate absorption probabilities of each cell towards one of the three defined terminal states: neural crest, neural and placodal. **(E)** Left: Gene module dynamics plots displaying GAMs of scaled gene module expression across latent time. GAMs are weighted by the fate absorption probability of each cell towards one of the three terminal states (placodal, neural crest and neural). Right: Heatmap of gene modules that display fate-specific expression (full list of gene modules available in Figure 4-Source Data 1). Gene modules were first unbiasedly filtered based on their differential expression between cell states (see methods and Figure 4-figure supplement 1A). Gene modules were further manually filtered based on expression patterns described in the literature (see results section: *Defining neural, neural crest and placodal gene modules*). Key genes of interest are highlighted on the right (for full gene list see Figure 4-Source Data 1).

When examining the genes within each module, we noticed that GM20 and GM22 include genes with widespread expression or expression in the lateral mesoderm and/or the extraembryonic region (*Epas1, Nkx2-5, Tnnc1, Pgk1*) (Adams et al., 2008; Bell et al., 2004; Liberatore et al., 2000; Ota et al., 2007) (Figure 4–Source Data 1). We therefore removed these modules from subsequent analysis. GM7 contains well characterised anterior markers including *Otx2, Six3* and *Pax6*, whilst GM11 contains midbrain/hindbrain markers including *Pax2*, *Nkx6-2* and *Wnt4* (Figure 4–Source Data 1). We find that at ss4 and ss8 these gene modules are indeed upregulated in the forebrain/aPPR and midbrain/hindbrain, respectively (Figure 4–figure supplement 1A). To focus our analysis on the segregation of neural, neural crest and placodal cells rather than on A-P patterning, A-P restricted modules were removed. The remaining five modules show broadly pan-neural (GM9), pan-placodal (GM21, GM23, GM24) or neural crest cell (GM13) expression at ss4 and ss8 (Figure 4E).

#### Latent time analysis suggests late emergence of neural crest cells

Given the cellular heterogeneity observed at early timepoints, discrete developmental stages do not necessarily accurately represent the developmental progression of individual cells towards a given fate. In contrast, latent time can measure the biological clock of individual cells as they differentiate and thus represents the order of cells along a developmental trajectory. To explore the expression dynamics of the five selected gene modules as neural, neural crest and placodal fates emerge, we first ordered cells across latent time using scVelo for the entire dataset (Figure 4C and Figure 4–figure supplement 1B). To investigate the segregation of all three fates, three separate trajectories need to be distinguished. To do this we calculated the probability of each cell to transit towards one of three terminal states (Figure 4–figure supplement 1C). These probabilities were obtained using CellRank (Lange et al., 2022), which leverages RNA velocity and transcriptomic similarity information to predict transitions between cells. Visualising the fate probabilities across the full dataset shows that cells with low latent time values (i.e., early cells) have low probabilities of becoming neural, neural crest or placodal cells (Figure 4D, see Figure 4A for stage reference). As cells ‘age’ the fate probability increases as they become transcriptionally similar to the terminal state. Unlike placodal and neural fates, neural crest fate probabilities are only high in ‘late’ cells (Figure 4D), reflecting the fact that defined neural crest cell clusters only appear at HH7.

#### Gene module dynamics reveals key differences in the activation of the transcriptional programmes associated to neural, neural crest and PPR specification

Having predicted the developmental trajectory of cells using scVelo and CellRank, we next modelled the expression of the five selected gene modules as a function of latent time with the aim to characterise their dynamics. Gene expression dynamics were modelled using generalised additive models (GAMs). GAMs were chosen because gene expression is highly dynamic and does not change linearly over time. For each gene module we fitted three separate GAMs, one for each fate (Figure 4E; left). The three fates were distinguished using the previously calculated fate absorption probabilities. This modelling approach reveals dynamic changes of gene expression. Three gene modules – GM9, GM13, GM23 – are initially inactive and are upregulated at different points across latent time in cells with neural, neural crest and placodal character, respectively.

GM9 becomes upregulated in all neural clusters and includes neural markers like *Lmo1* and *Sox21* (Figure 4E ‘Neural GM9’, Figure 4–Source Data 1). Although the expression of different genes within this module correlates broadly, there is some gene-gene variation. For example, *Lmo1* is the first gene to be expressed and exhibits highly dynamic expression over time. It is first found at HH5 within the eCN, pNP and iNP, before being upregulated across the entire neural plate at HH7 and ss4. At ss8, *Lmo1* is specifically downregulated within the MB. In contrast, *Sox21* is first upregulated at HH5/6 and is consistently expressed throughout all neural cell clusters.

In contrast, modelling reveals that GM13 is upregulated in cells with neural crest identity. Indeed, it contains well known neural crest markers *Pax7, Snai2, Sox10* and *Tfap2b* (Figure 4E ‘NC GM13’, Figure 4–Source Data 1) and is initially broadly activated in NPB and neural crest clusters at HH7. At ss4 and ss8, this module remains strongly expressed within neural crest and delaminating neural crest clusters. Within GM13 individual genes are activated at different time points: *Pax7* is the first gene to be expressed in the pNPB at HH5, while *Sox10* starts to be expressed only from ss4 onwards.

Finally, modelling the expression dynamics of GM23 shows that it is activated early and becomes upregulated in cells with placodal character across latent time. This prediction is supported by the fact that GM23 is highly expressed in PPR clusters at ss8 and contains *bona fide* placodal genes like *Six1* and *Eya2* (Figure 4E ‘Placodal GM23’, Figure 4–Source Data 1). At HH4, most GM23 genes are not expressed, but are broadly activated at HH5 in PPR, non-neural and NPB clusters, where they remain active until becoming restricted to PPR cells at ss4. Comparing the onset of activation of these three gene modules across latent time reveals a clear temporal order with the placodal module being activated first, followed by the neural and neural crest gene modules.

In contrast to these modules, modelling of GM24 and GM21 shows that they are initially expressed in all three cell populations before being rapidly downregulated within neural crest and neural cells (Figure 4E ‘Epiblast GM21’ and ‘Epiblast GM24’). Indeed, genes in GM24 are broadly expressed across the epiblast at HH4, and then become gradually restricted to PPR, non-neural and NPB clusters at HH6/7, and to PPR cells at ss4/8 (Figure 4A). GM21 displays similar dynamics but is restricted to non-neural clusters earlier at HH5. Although some genes within these epiblast modules have been described as non-neural/PPR specific based on *in situ* hybridisation (*Tfap2c, Bambi, Homer2, Tspan13*) (Bell et al., 2004; Hintze et al., 2017; Mehdizadeh et al., 2021; Reichert et al., 2013; Rothstein & Simoes-Costa, 2020), others are found to be expressed within the early neural territory (*Epcam*, *Sall4*, *Cited4*) (Barembaum & Bronner-Fraser, 2007; Bell et al., 2004; Trevers et al., 2021) (Figure 4–Source Data 1).

In summary, modelling the expression dynamics of selected gene modules provides new insight into the sequential activation of fate-specific developmental programmes, as well as proposing key differences in the mechanism by which neural, neural crest and placodal cells are specified. We show, using an unbiased gene module approach, that a placodal gene module is activated first, followed by neural and neural crest modules. While neural and neural crest modules are specifically activated in the relevant cell clusters alone, two modules that are later restricted to the PPR are initially broadly expressed across the epiblast. We therefore propose that the specification of neural and neural crest states may require the downregulation of epiblast modules which later become confined to placodal cells.

### Spatially restricted co-expression of genes at the NPB

Previously, the NPB has been defined based on the expression of so-called NPB specifiers, however our transcriptome analysis suggests that these markers have either a placodal or neural crest bias (Figure 3D). The question remains as to where NPB specifiers are co-expressed; are they co-expressed throughout the border, or are putative unstable progenitors spatially restricted? To investigate this, we sought to model the expression of ‘NPB specifiers’ across the mediolateral axis, before validating their spatial expression *in vivo* using *in situ* hybridisation chain reaction (HCR).

We chose to model the spatial expression of ‘NPB specifiers’ using our HH7 scRNAseq data as this stage marks the start of placodal and neural crest segregation (Figure 3A). Principal component analysis reveals that HH7 cells are ordered along the inverse of principal component (PC) 1 based on their M-L position *in vivo*; PC1 therefore provides an axis through which we can model spatial gene expression (Figure 5A). Smoothed expression profiles across PC1 were modelled using GAMs calculated for each NPB specifier (Figure 5B).

**Figure 5.**
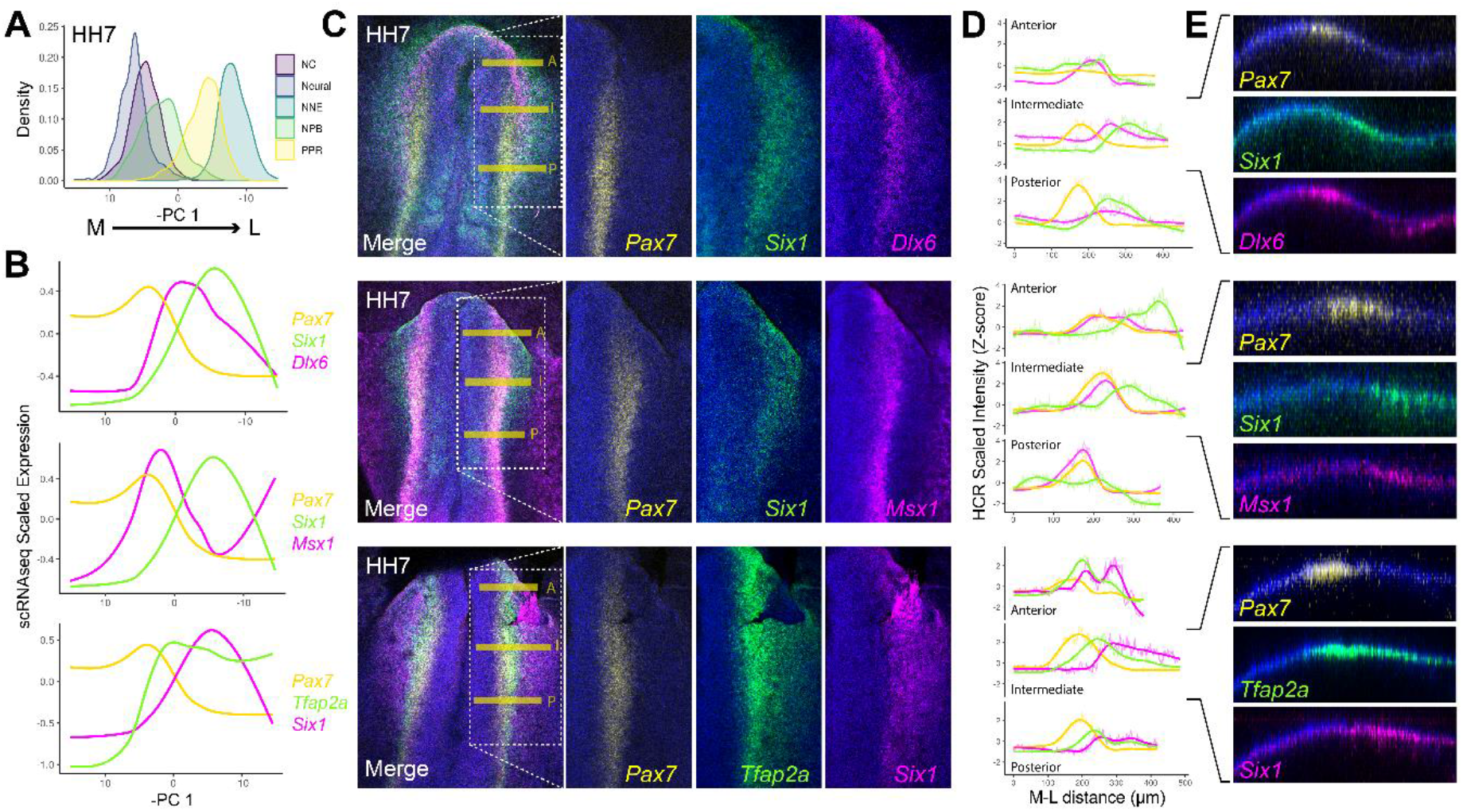
HCR validates spatial restriction of classical NPB specifiers. **(A)** Distribution of cells at HH7 from major ectodermal cell lineages across the inverse of principle component 1 (PC 1), revealing medio-lateral patterning across this axis. **(B)** Spatial gene expression modelling of key placodal and neural crest specifiers at HH7 across the inverse of PC 1. GAMs were fitted for each gene to predict their medio-lateral pattern of expression in the embryo. **(C)** Whole mount *in situ* HCR images at HH7 for combinations of markers modelled in B. Overlayed dotted boxes in the merged image show the region of interest displayed for each separated colour channel; yellow lines indicate the line where intensity measures in D were taken **(D)** Fluorescent intensity measurements taken at anterior ‘A’, intermediate ‘I’ and posterior ‘P’ regions depicted by the yellow bars in C. Intensity measurements were scaled for each gene across the three axial levels to allow for relative comparisons between different axial regions. **(E)** Virtual cross sections taken from the intermediate region, highlighting expression of each marker within the embryonic ectoderm.

Our models predict that *Msx1* and *Pax7* are upregulated medially relative to *Dlx6* and *Six1*, in line with their roles in neural crest and placodal specification, respectively (Basch et al., 2006; Brugmann et al., 2004; Christophorou et al., 2009; McLarren et al., 2003; Monsoro-Burq et al., 2005). Interestingly, *Pax7* and *Six1*, which are upregulated later than either *Msx1* and *Dlx6* (Figure 3E), are predicted to exhibit a smaller region of co-expression with each other than with either *Msx1* or *Dlx6* (Figure 5C). This highlights that *Pax7* and *Six1* have a significant bias towards either neural crest/placodal fates. *Tfap2a*, which is required for both placodal and neural crest specification (Bhat et al., 2013; Rothstein & Simoes-Costa, 2020), is predicted to overlap extensively with both *Pax7* and *Six1*.

HCR at the same stage validates the predicted levels of co-expression across the M-L axis *in vivo* (Figure 5C-E). We quantified M-L expression by taking intensity measurements at three different A-P positions (Figure 5C; yellow horizontal lines). The intensity measurements for each gene were Z-scored across all three axial levels to allow for comparative relative measurements of gene expression (Figure 5D). We note a striking correlation in the expression intensity measurements taken from the intermediate axial level (Figure 5D; Intermediate) and the predicted M-L expression patterns from our scRNAseq data (Figure 5B), which represent an aggregate of all A-P regions.

Taken together, both our scRNAseq and HCR analysis reveal segregation of neural crest and placodal markers across the M-L axis, with considerable heterogeneity in gene co-expression depending upon the NPB markers in question. Importantly, there are striking differences of gene expression and co-expression along the A-P axis. While *Dlx6* and *Six1* are strongly expressed anteriorly, *Pax7* and *Msx1* show increased expression levels posteriorly (Figure 5D). Thus, cells within the anterior NPB may be biased towards placodal fates, whereas cells located posteriorly may be biased towards neural crest cells. These findings suggest that only a small subset of cells at the NPB are in an unstable state. Furthermore, while the NPB can be defined as a territory adjacent to the developing neural plate, a unique NPB cell state coinciding with this territory cannot be defined transcriptionally using previously established criteria.

### BLUPs: multi-potent progenitors at the NPB?

Given the cell heterogeneity at the NPB, we sought to define an approach that did not depend on the expression of definitive NPB specifiers to identify and label transcriptionally unstable progenitors. Transcription factors that specify neural crest and placodal fates are known to cross-repress one another whilst enhancing their own expression (for review: Thiery et al., 2020). Therefore, cells that co-express placodal and neural crest genes may represent unstable progenitors which retain plasticity; we term these co-expressing cells ‘Border-Located Unstable Progenitors’ (BLUPs).

#### Defining neural crest and placode specific gene modules

To locate BLUPs within the ectoderm, we identified gene modules that are restricted to either neural crest or placodal cells at late stages (ss8) and investigated their co-expression at earlier stages. 15 modules were differentially expressed in at least one cell state (Figure 6–figure supplement 1A and Figure 6–Source Data 1). GM9-13 and GM16-17 are characteristic for neural cells with different A-P identities. Seven gene modules are absent from neural populations but expressed in placodal cells (GM4-6), neural crest (GM1-2) or both (GM7-8). GM5 contains well characterised placodal markers including *Six1, Homer2* and *Znf385c* and is expressed in both aPPR and pPPR clusters. This module was therefore selected as a pan-placodal module. GM2 is expressed in both neural crest and delaminating neural crest clusters and contains neural crest specifiers like *FoxD3, Snai2, Sox9, Pax7* and *Msx1*, as well as the Wnt ligands *Wnt1* and *Wnt6*. GM2 thus represents a *bona fide* neural crest specific module. Although GM1 contains multiple neural crest markers (*Ets1, Sox10, Sox8*; Figure 6–Source Data 1), it is only expressed in delaminating neural crest cells and was therefore excluded for further analysis. Thus, GM5 and GM2 were considered as pan-placodal and pan-neural crest module and used for co-expression analysis (Figure 6A).

**Figure 6.**
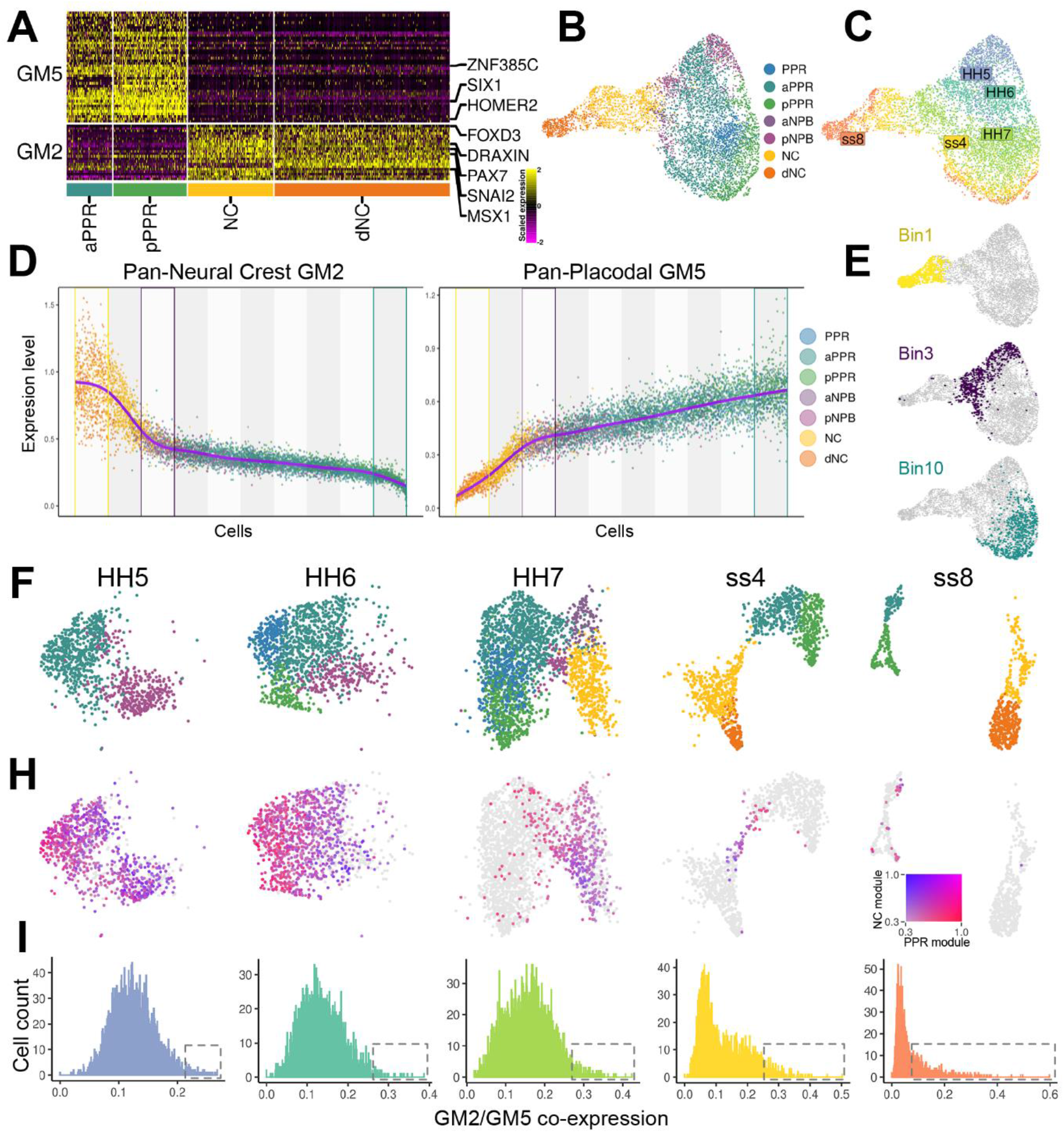
Border-Located Unstable Progenitors’ (BLUPs) co-express placodal and neural crest gene modules. **(A)** Heatmap displaying pan-placodal (GM5) and pan-neural crest (GM2) at ss8 which have been subset from gene modules in Figure 6-figure supplement 1A. **(B)** UMAP plot of the NPB subset coloured by cell state. **(C)** UMAP plot of the NPB subset coloured by developmental stage. **(D)** Average normalized expression of placodal and neural crest gene modules (GM5 and GM2, respectively) identified at ss8 (see Figure 5). Cells are ordered by their expression ratio of these two modules and coloured by cell state. Cells were grouped into 10 evenly sized bins (background shaded regions). Three bins (1, 3 and 10) are overlaid to highlight cells with high neural crest/low placodal (bin 1), equivalent levels of both (bin 3) and high placodal/low neural crest (bin 10) gene expression. **(E)** UMAP plots of the NPB subset displaying the cells captured within bins 1, 3 and 10. **(F-H)** UMAP plots from each developmental stage, displaying only cells in the NPB subset (equivalent to red cells in Figure 6-figure supplement 1B). **(F)** Cells are coloured by cell state. **(H)** Co-expression analysis of the pan-neural crest and pan-placodal modules (see methods). Cells which co-express both modules above 0.3 are coloured. **(I)** Histograms revealing the distribution of co-expression of the pan-neural crest and pan-placodal modules at each developmental stage. At later stages (ss4-ss8), the distribution of co-expression shifts from a normal to a negative binomial distribution.

### BLUPs co-express placodal and neural crest lineage specifiers

To investigate the co-expression of these modules in NPB cells and their derivatives, we first selected NPB (aNPB, pNPB), neural crest (NC, dNC) and PPR (aPPR, pPPR, PPR) cell clusters from HH5-ss8 (Figure 6–figure supplement 1B, red cells; thereafter referred to as NPB subset). Next, we re-clustered this subset of cells (Figure 5B). UMAP analysis highlights a bifurcation of neural crest and placodal fates between HH7 and ss4, with the NPB located at the boundary between neural crest and aPPR cell clusters (Figure 6B, C).

To examine the relative expression of pan-placodal and pan-neural crest modules within the NPB subset, we ordered cells according to the expression ratio of both modules and grouped them into 10 evenly sized bins (Figure 6D and Figure 6–figure supplement 2). Cells at the extreme ends exclusively express the pan-neural crest (Bin 1) or the pan-placodal gene module (Bin 10). As expected, these cells represent mostly delaminating neural crest or placodal cells, respectively. However, cells in Bin 3 show equal expression levels of both modules and contain a mixture of NPB, PPR and neural crest cells (Figure 6D). Mapping Bin 3 cells back to the UMAP revealed that BLUPs are located specifically between PPR and neural crest cell clusters (Figure 6E).

To evaluate the distribution of BLUPs at different developmental timepoints, we mapped Bin 3 cells back to the UMAPs of individual stages (Figure 6–figure supplement 1C). We find at HH5 and HH6, Bin 3 cells map to aPPR and pNPB clusters, whilst they map predominantly to neural crest and NPB clusters at HH7. At ss4, there is a sharp reduction in the number of Bin 3 cells. Interestingly, although there is no cluster classified as NPB at this stage, BLUPs are still located at the boundary between neural crest and PPR clusters and are predicted to be in an undetermined transcriptional state by RNA velocity (Figure 3G). Once placodal and neural crest fates are largely segregated at ss8, only very few scattered cells express similar levels of pan-neural crest and pan-placodal modules.

The analysis above focuses on cells expressing similar levels of pan-placodal and pan-neural crest modules. Next, we visualised cells that co-express these modules at different levels. We calculated pan-neural crest and pan-placodal module co-expression per developmental stage and filtered cells below a minimum heuristic threshold of 0.3 (Figure 6H). At HH5/6 almost all cells classified as NPB and placodal cells co-express both modules above the threshold. However, this changes at HH7, when co-expressing BLUPs are largely restricted to the NPB and neural crest clusters. By ss4, only a small population of BLUPs found between placodal and neural crest clusters continue to co-express alternative fate markers. Over time, we note a striking shift in the distribution of co-expression between cells (Figure 6I). At early stages (HH5-7) most cells co-express pan-placodal and pan-neural crest gene modules at similar levels, whilst at later stages (ss4 and ss8) only a small subset of cells retain high relative levels of co-expression (Figure 6I, boxes). This shift in distribution in the number of highly co-expressing cells suggests that placodal/neural crest segregation takes place from around HH7 until ss8.

This analysis highlights that BLUPs are transcriptionally unstable, co-expressing gene modules that characterise both neural crest and placodal cells. This expression profile is likely to endow them with the potential to give rise to both, if not all NPB fates. The NPB demarcates an anatomical region surrounding the neural plate, however it lacks a definitive transcriptional signature highlighting that it is transcriptionally heterogeneous. Given the previously characterised cross repressive interactions between placodal and neural crest specifiers (for review: Thiery et al., 2020), we propose that unstable NPB progenitors (BLUPs) are better defined based on the co-expression of alternate gene modules. The proportion of these cells decreases over time, but a small population at ss4 retains this characteristic and is therefore likely to retain bi- or multipotency even as the neural tube closes. We have shown that BLUPs are likely in an unstable transcriptional state, however how this transcriptional heterogeneity is resolved requires further exploration.

### Identifying a temporal gene expression hierarchy regulating the specification of neural crest and placodal fates

To investigate how BLUPs segregate towards neural crest and placodal fates, we modelled gene module dynamics across the NPB subset (for workflow explanation see Figure 4–figure supplement 2). We previously characterised gene expression dynamics (Figure 4E) across the whole dataset and uncovered sequential activation of placodal, neural and neural crest gene programmes. However, focusing on the NPB subset alone provides a higher resolution of the gene expression hierarchy regulating the segregation of placodes and neural crest.

To model gene expression dynamics during lineage segregation at the NPB, we performed scVelo analysis on the NPB subset (Figure 7A) and used CellRank to determine the probability that each cell will become neural crest or placodal (Figure 7A-D). We then identified putative lineage specifiers by calculating gene modules which are differentially expressed between placodes and the neural crest. These 10 differentially expressed gene modules were further filtered to include only those that contained genes expressed in neural crest, delaminating neural crest or PPR cell states (as defined in the binary knowledge matrix). This analysis identified three modules (GM12-14) upregulated in placodal and four in neural crest cells (GM40 and GM42-44) between HH7 and ss8 (Figure 7E, Figure 7–Source Data 1). Using our latent time and lineage probability measurements, we model the dynamics of these 7 gene modules during the segregation of the placodes and neural crest (Figure 7E; left).

**Figure 7.**
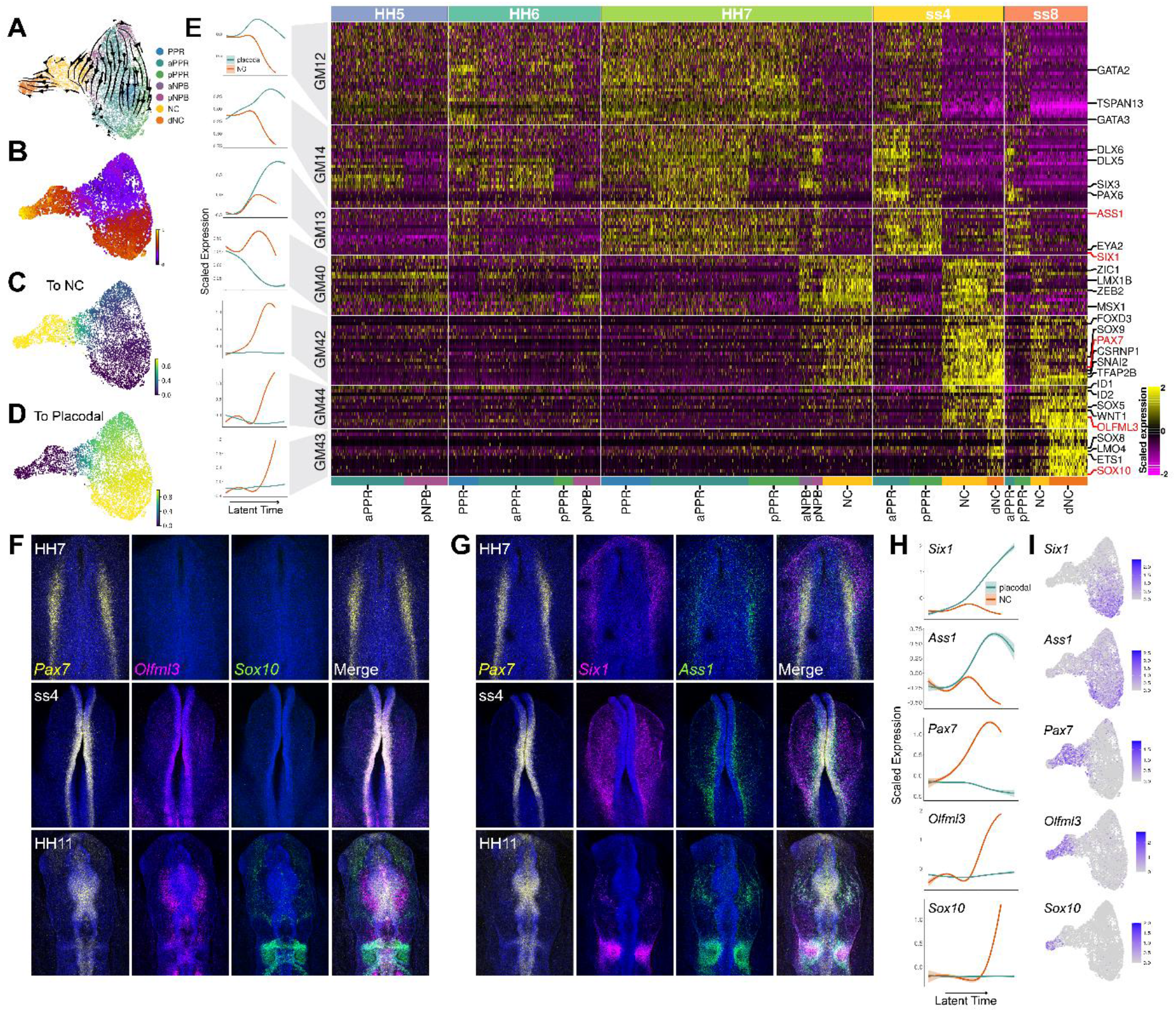
NPB gene module analysis reveals temporal hierarchy of gene expression during the specification of neural crest and placodes. **(A)** UMAP plot with cells coloured by cell state classifications and overlaid with RNA velocity vectors depicting the predicted directionality of transcriptional change. **(B)** UMAP plot of the NPB subset with cells coloured by latent time. **(C-D)** UMAP plots showing the fate absorption probabilities of each cell towards the specified terminal states (neural crest or placodes). **(E)** Left: Gene module dynamics plots displaying GAMs of scaled gene module expression across latent time. GAMs are weighted by the fate absorption probability of each cell towards either placodal or neural crest terminal states. Right: Heatmap showing gene modules calculated across the NPB subset (full list of gene modules available in Figure 7-Source Data 1). To identify gene modules potentially involved in placodal or neural crest specification, gene modules were filtered to include those that show differential expression between placodal and neural crest cell states (see methods). Genes specified as neural crest or placodal in our binary knowledge matrix (Supplementary File 1) were selected as bait genes to further filter the gene module list for visualisation. Known and novel placodal and neural crest markers highlighted in red were validated by *in situ* HCR (F-G). **(F)** Whole mount *in situ* HCR at HH7, ss4 and HH11 validating the expression of *Olfml3* at the NPB and delaminating neural crest. **(G)** Whole mount *in situ* HCR at HH7, ss4 and HH11 validating the expression of *Ass1* in the pre-placodal region. **(H)** Gene dynamics displaying GAMs of scaled gene expression across latent time. GAMs are weighted by the fate absorption probability of each cell towards either placodal or neural crest lineages. **(I)** Feature plots for genes validated by *in situ* HCR on the HH7 stage UMAP.

Of the placodal modules, GM14 and GM12 are expressed as early as HH5. GM12 is expressed throughout both aPPR and pNPB and contains early non-neural/PPR markers like *Gata2/3* (Figure 7E, Figure 7–Source Data 1). At HH6, GM12 is downregulated in the NPB and remains confined to the PPR clusters until ss8. Unlike GM12, GM14 is initially restricted to the PPR and only activated in the NPB at HH7. At ss4 and ss8 its expression is confined to the PPR, with a higher expression in anterior cells. This module contains pan-placodal genes *Dlx5/6* as well as anterior markers *Six3* and *Pax6*. GM13 is the last placodal module to be activated and contains the definitive placodal transcripts *Six1* and *Eya2*. It is first expressed at low levels at HH6 before being broadly activated across the PPR at HH7 through to ss8. We also observe weaker expression of this module at the NPB cells at HH7. To visualise the sequential activation of these modules across latent time, gene expression of each cell was weighted based on its placodal fate probability. This analysis shows that GM12 is activated before GM14 and GM13 in latent time. Whilst the expression of GM12 and GM14 does not change dramatically, GM13 is highly upregulated over latent time within the placodal cells (Figure 7E and Figure 7–figure supplement 1A).

The four neural crest gene modules each contain *bona fide* neural crest transcriptional regulators including *Msx1, Zic1, Zeb2, Lmx1b* (GM40), *Pax7, Snai2, Axud1, Foxd3, Tfap2b, Sox9* (GM42), *Sox5, Id1/2* (GM44) and *Sox10, Ets1, Lmo4* and *Sox8* (GM43) (Figure 7E and Figure 7–Source Data 1). Our analysis groups these factors unbiasedly into four modules; modelling their expression across latent time weighted by neural crest fate probability shows a clear sequential activation from GM40 to GM42, GM44 and GM43 (Figure 7-figure supplement 1). GM40 is activated first, initially broadly across aPPR and pNPB clusters at HH5 and HH6, before becoming restricted to neural crest and NPB clusters from HH7 onwards. By ss8, it is downregulated in the delaminating neural crest population. GM42 and GM44 are highly upregulated only in neural crest and delaminating neural crest cells, with GM42 first expressed at HH7 and GM44 at ss4. Lastly, GM44 is activated specifically within the delaminating neural crest cells at ss4 where it is further upregulated at ss8 (Figure 7E). This order suggests a hierarchical relationship between these factors in the neural crest gene regulatory network.

While gene module selection was based on prior knowledge of marker genes, this analysis also allows us to predict novel factors with similar expression profiles. We selected *Ass1* and *Olfml3* to validate the expression of novel placodal and neural crest markers, respectively, using HCR (Figure 7F, G). Like the placodal specifiers *Six1* and *Eya1, Ass1* is found in GM13. HCR confirms that *Ass1* and *Six1* are co-localised within the lateral NPB at HH7 and ss4, and in the otic placode at HH11 (Figure 7G). However, we also note a greater degree in co-expression of *Pax7/Ass1* than *Pax7/Six1* (Figure 7G; HH7 and ss4), as expected given our gene expression dynamics predicts that *Ass1* is initially upregulated in both placodal and early neural crest cells before it is downregulated in the neural crest (Figure 7H, I). *Olfml3* was chosen as a novel neural crest marker, as it is found in GM44 which is predicted to be upregulated within the neural crest in between GM42 (Pax7) and GM43 (*Sox10*) (Figure 7H). HCR confirmed the predicted sequential temporal expression of these markers, with *Pax7* active in the posterior NPB at HH7, *Pax7* and *Olfml3* co-expressed at NPB at ss4, and all three markers expressed within the neural crest at HH11 when the neural crest is undergoing delamination (Figure 7F).

In summary, analysing gene module dynamics across the NPB subset reveals the sequential activation of different co-regulated genes in placodal and neural crest cells. Many genes within these modules have been previously characterised and are known to form part of the neural crest or placode gene regulatory network (for review: Grocott et al., 2012; Pla & Monsoro-Burq, 2018; Schlosser, 2006, 2014; Simoes-Costa & Bronner, 2015; Thiery et al., 2020). This unbiased approach groups these factors and predicts a hierarchical order, as well as identifying new candidate factors which may play important roles in the segregation of the neural crest and placodal fates.

## Discussion

The regulation of cell fate specification at the NPB has been extensively studied in a range of model systems, giving rise to a wealth of knowledge regarding transcriptional regulators and their interactions. Nevertheless, our understanding as to how individual cells undergo cell fate decisions has been limited by our inability to study transcription and gene regulation at a single cell level. Single cell immunohistochemistry has demonstrated that in contrast to what was previously thought, cells co-express markers that specify either placodal, neural or neural crest fates, even at late neurulation (Roellig et al., 2017). A recent single cell RNAseq analysis has attempted to characterise transcriptional heterogeneity in the ectoderm, and proposed that a definitive NPB is not established until early neurulation (HH7), at which point placodal and neural crest lineages gradually segregate (Williams et al., 2022). However, an absence of definitive placodal cells collected at late neurulation, as well as a low number of ectodermal cells sampled limits the overall characterisation of ectodermal heterogeneity and makes conclusions about placodal and neural crest segregation problematic. Nevertheless, newly characterised cellular heterogeneity requires us to revisit models for cell fate specification at the NPB and to reconsider our definition of the NPB itself.

In this study, we characterise ectodermal cell states and cellular heterogeneity from primitive streak stages though to late neurulation in chick. We demonstrate that although significant transcriptional differences are apparent between cells at primitive streak stages, due to high levels of heterogeneity, cells do not cluster into discrete cell states exhibiting restricted expression patterns until early neurulation (HH7). Instead, cell states are initially characterised by the overlapping expression of broad early neural and non-neural genes, as well as axial markers. We identify increased cell state diversification during neurulation, and segregation of neural, neural crest and placodal lineages taking place across 1-somite to 8-somite stages. High resolution transcriptional analysis of cells at the NPB reveals that by the 4-somite stage, placodal and neural crest lineages have largely segregated. However, despite this segregation, we continue to identify high levels of co-expression of placodal and neural crest genes within BLUPs which are predicted to be in a dynamic transcriptional state.

### Specification of neural and neural crest cell lineages requires downregulation of broad epiblast modules

Although specification networks for the neural, neural crest and placodal lineages are well established (for review: Betancur et al., 2010; Grocott et al., 2012; Pla & Monsoro-Burq, 2018; Stundl et al., 2021; Thiery et al., 2020), the order in which these lineages are specified remains unclear. Our unbiased gene module analysis has identified groups of genes which correlate and are specifically upregulated within each cell fate. Our gene modules support the specification networks established in previous studies. *Pax7, Tfap2b, Snai2* and *Sox10* cluster together and are upregulated in neural crest states, while *Six1, Eya2, Gata2/3* and *Dlx5/6* are activated within placodal states and *Lmo1* and *Sox21* in neural states.

We also find putative novel regulators of these cell states that are co-regulated with known fate specifiers, as well as the order in which these cell fates are specified. Placodal, neural and neural crest specific gene modules are activated sequentially, suggesting that these states are specified in that order. However, there is also asynchrony in the initiation of cell lineage specification; although we classify placodal cells as early as HH5, we also identify transcriptionally unstable BLUPs at ss4. This suggests cell fate allocation at the NPB is a gradual process, with the number of unspecified progenitors decreasing over time as they commit to a given lineage.

Alongside characterising placodal, neural and neural crest gene modules, we observe the expression of two modules which exhibit initial broad pan-epiblast expression at primitive streak stages, but which are confined to placodal cell states at the 4 and 8-somite stages. Within these epiblast modules, unsurprisingly we note the expression of important placodal and non-neural genes *Tfap2c, Homer2* and the BMP inhibitor *Bambi*. In addition, we also find the pluripotency marker *Sall4* and *Tspan13*, with the latter shown to be positively regulated by Six1 (Mehdizadeh et al., 2021). A recent study in chick suggests that based on transcriptional profiling a ‘pre-border’ state can already be defined prior to gastrulation. *In vitro* experiments show that the pre-gastrula epiblast is specified as NPB; once cells have acquired a ‘pre-border’ state they can become neural and placodal precursors depending on the signalling input (Trevers et al., 2018). Given that the expression of epiblast modules becomes confined to the placodes, we speculate that the specification of the neural crest and neural states might require the downregulation of these modules.

### Identifying BLUPs at the neural plate border based on the co-expression of competing gene modules

Members of Zic, Tfap2, Pax, Msx and Dlx transcription factor families have previously been termed NPB specifiers (for review: Stundl et al., 2021; Thiery et al., 2020). Tfap2a has been identified as a pioneer factor which triggers the expression of other NPB specifiers and is required but not sufficient for both neural crest and placodal cell fates (Bhat et al., 2013; Rothstein & Simoes-Costa, 2020). The cell fate switching role of Tfap2a depends upon its association with its co-factors, Tfap2b and Tfap2c. Tfap2c is necessary for the induction of NPB specifiers *Msx1, Pax7* and *Zic1*, whilst Tfap2b is later required for downstream neural crest specification (Rothstein & Simoes-Costa, 2020).

Gene module analysis across the neural plate border highlights the sequential activation and restriction of previously characterised NPB specifiers to neural crest cells. Early NPB specifiers *Msx1* and *Zic1* are initially broadly expressed across pNPB and aPPR cells at HH5, but later become downregulated in placodal cells and upregulated in the neural crest. The gene module containing *Pax7* is activated slightly later and contains other NPB specifiers *Axud1* and *Tfap2b*, as well as definitive neural crest markers *FoxD3* and *Snai2*. Unlike the *Msx1* module, which is initially upregulated in both aPPR and pNPB clusters at HH5, the *Pax7* module is specifically restricted to the NPB and subsequent neural crest clusters (Figure 7E). This analysis reveals that despite these markers having been previously characterised as NPB specifiers, their expression becomes restricted to the neural crest. In contrast, *Dlx5/6* and *Tfap2a* are both found within a gene module that is later restricted to the placodes (Figure 4E).

Although *Tfap2a* is grouped in the same module as placodal specifiers *Six1* and *Eya2*, it is expressed in both placodal and neural crest cell states, even at later stages (Figure 3D, E). This expression reflects the key role of *Tfap2a* as a pioneer factor in both neural crest and placodal specification, however its expression alone is not sufficient to classify putative multi-potent progenitors at the NPB. At HH5, although *Tfap2a* is expressed within the pNPB cluster alongside other NPB specifiers, it is also expressed within aPPR cells which never express NPB specifiers including *Pax7, Pax3* and *Tfap2b* (Figure 2E).

To classify the NPB as a discrete state, NPB markers need to be uniformly upregulated in NPB cells which give rise to neural, non-neural ectoderm, neural crest and PPR fates. Given the non-uniform expression of previously characterised NPB specifiers (Basch et al., 2006; Ezin et al., 2009; Hong & Saint-Jeannet, 2007; McLarren et al., 2003; Rothstein & Simoes-Costa, 2020; Streit & Stern, 1999), and the expression of *Tfap2a* in the early aPPR cells which are unlikely to give rise to other fates, we propose that the NPB cannot be accurately classified using just these previously established genes. Instead, the NPB is an anatomical region surrounding the neural plate within which putative unstable progenitors (BLUPs) reside. BLUPs can be identified through the co-expression of gene modules which regulate the specification of alternative cell fates.

### Gradient border model resolves the neural plate border and binary competence dilemma

The ‘binary competence’ and ‘neural plate border’ models have provided useful frameworks for understanding cell fate determination during neural, neural crest and placodal segregation (Figure 8A, B) (for review: Schlosser, 2006, 2014; Thiery et al., 2020). The binary competence model suggests that the ectoderm is subdivided into non-neural/placodal and neural/neural crest domains (Ahrens & Schlosser, 2005; Pieper et al., 2012), whilst the neural plate border model suggests cells at the border remain multi-potent and can differentiate into all ectodermal fates (Baker & Bronner-Fraser, 1997b; Roellig et al., 2017; Streit & Stern, 1999). Due to the limitations of studying cell fate competence at a population level, transplantation experiments have failed to resolve these two models.

**Figure 8.**
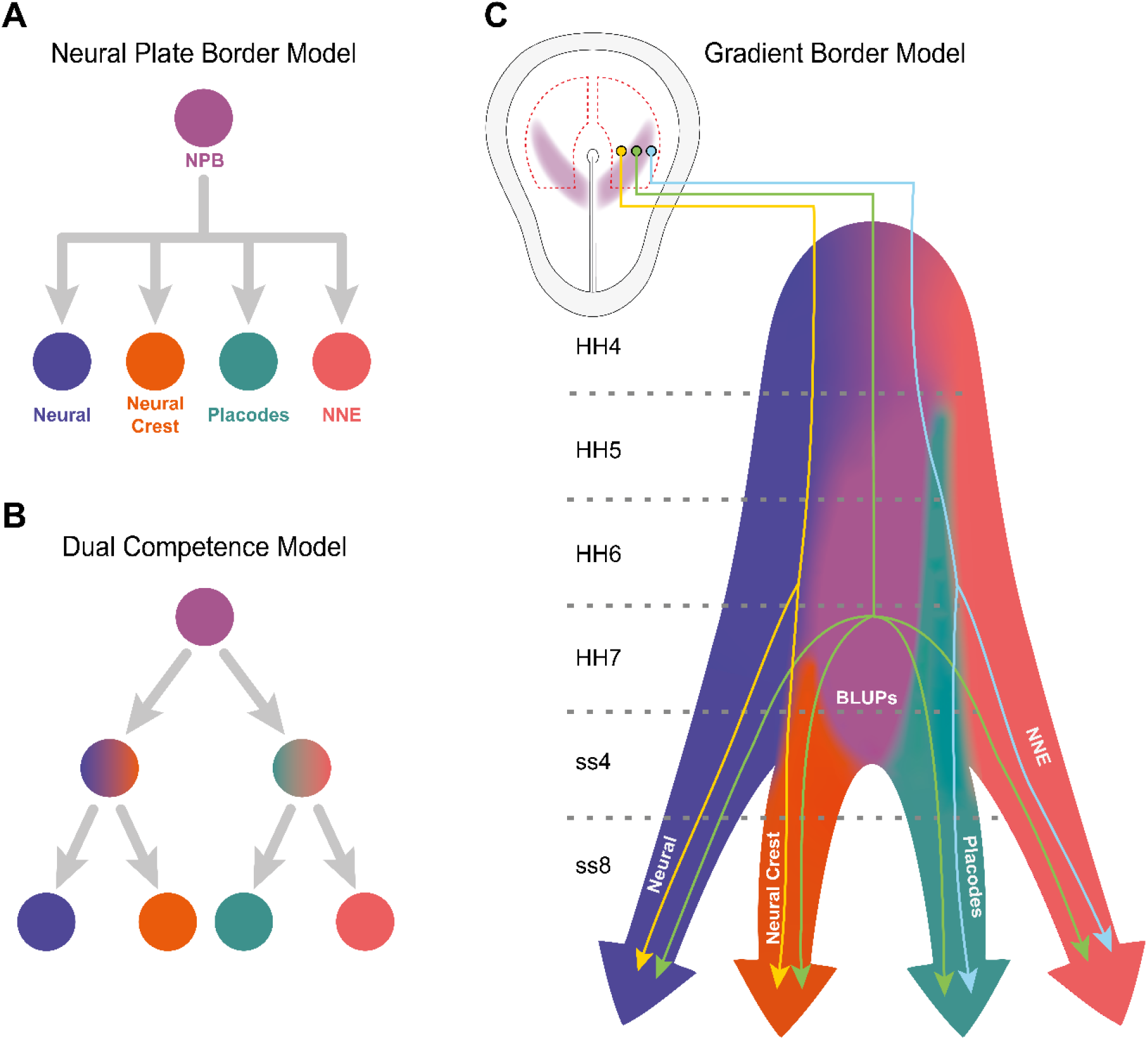
Gradient border model resolves conflicting models of lineage segregation at the neural plate border. **(A)** Schematic illustrating the neural plate border model. This model illustrates that cells at the neural plate border are multipotent and able to give rise to neural, neural crest, placodal and non-neural ectoderm (NNE) lineages. **(B)** Schematic illustrating the dual competence model. This model suggests that the neural plate border is first defined into non-neural/placodal and neural crest/neural competence domains prior to further subdivision into each of the final four cell lineages. **(C)** Schematic illustrating the gradient border model. This model proposes a probabilistic model of cell fate allocation which is intrinsically linked to the spatiotemporal positioning of a cell. We suggest that cells located within the medial neural plate border (yellow cell lineage) will give rise to neural and neural crest, cells within the lateral neural plate border (blue cell lineage) to non-neural ectoderm and placodes, whilst subsets of cells (including BLUPs) will remain in an indeterminate state and continue to give rise to all four lineages even at late stages of neurulation (green cell lineage). Shaded background colours represent the cell state space.

Our co-expression analysis of pan-neural crest and pan-placodal gene modules revealed that most cells in the early NPB can be classified as BLUPs, as they co-express these modules at similar levels. At later stages, we see a divergence in the level of co-expression between cell populations, with the highest co-expressing cells located at the branching points between the neural crest and placodal lineages (Figure 6H and I). In summary, we find that over time, the BLUP population reduces in size until it is restricted to only a small number of cells at the boundary between placodal and neural crest cell clusters at the 4-somite stage. Cell fate allocation within BLUPs is likely to be driven via transcription factor cross-repression, given the previously characterised cross-repressive interactions taking place between placodal, neural crest and neural specifiers (Brugmann et al., 2004; Christophorou et al., 2009; Schlosser et al., 2008; Wakamatsu et al., 2004).

Following these findings, we propose a ‘gradient border’ model for cell fate allocation (Figure 8C). This model suggests that cell fate allocation is probabilistic and intimately linked to a cells spatiotemporal positioning. In accordance with the binary competence model, cells located within the lateral NPB will give rise to epidermal and placodal fates, whilst those found medially give rise to neural and neural crest fates. However, instead of a binary segregation of the NPB into neural/neural crest and non-neural/placodal domains, we suggest that BLUPs at the NPB are kept in an indeterminate state whilst they continue to co-express factors involved in the specification of alternative lineages. Both our co-expression analysis and previous documentation of heterogeneity at the NPB support this model.

## Materials and Methods

### Data collection and alignment

#### Single cell dissociation and 10X single-cell mRNA sequencing

Fertilised chicken eggs were incubated at 38°C for 24-32 hours depending upon the developmental stage collected. Embryos were pinned and had the endoderm and mesoderm removed using dispase (10mg/ml) before dissecting the ectodermal region of interest (Figure 2A). Multiple embryos from the same stage were pooled prior to dissociation. Samples were then dissociated in 250μl pre-heated FACSmax cell dissociation solution (Amsbio, T200100) with papain (30U/ml; Sigma, P3125) for 20 minutes at 37°C. Samples were gently pipetted every 5 minutes to facilitate dissociation. After dissociation, 250μl resuspension solution was added to prevent any further dissociation (HBSS; non-acetylated BSA (1mg/ml; Invitrogen, 10743447); HEPES (0.01M); non-essential amino acids (1x; Thermo Fisher Scientific, 11140050); rock inhibitor Y-27632 (10μM; Stemcell Technologies). Cells were passed through a 20μm cell strainer (Miltenyi Biotech, 130-101-812) to remove any debris, before pelleting and resuspending in 500μl resuspension buffer. To remove any dead or dying cells as well as any remaining doublets, 1μl 0.1mg/ml DAPI was added to the cell suspension before FAC sorting. Given the low number of cells obtained from each embryo, samples for each stage were collected over multiple days and stored in 90% MeOH with 10% resuspension solution after FAC sorting. Prior to sequencing, samples from the same stage were pooled and resuspended in DPBS with 0.5% non-acetylated BSA and 0.5U/μl RNAse inhibitor (Roche 3335399001).

Cells underwent single-cell mRNA sequencing at the Francis Crick Institute, London. Library preparation was carried out using the 10x 3’ single-cell mRNA v3 kit (10x Genomics), before sequencing on the HiSeq 4000 (Illumina) to a target depth of 50k reads/cell. Samples were sequenced in two batches (batch 1: HH4, HH6, ss4, ss8; batch 2: HH5, HH6, HH7, ss4), with two stages (HH6 and ss4) sequenced in both batches to allow for the correction of any potential batch effects. Data from the first batch has previously been used for the characterisation of neural induction (Trevers et al., 2021) (ArrayExpress accession number: E-MTAB-10408).

#### Read alignment

A custom pipeline was developed in Nextflow (v20.07.1) (Di Tommaso et al., 2017) to handle all GTF processing and read alignment whilst maintaining full reproducibility.

Prior to scRNAseq alignment, the GalGal6 ensembl 97 GTF file was modified to prefix genes from the Z and W sex chromosomes with their respective chromosome IDs (Z- and W-). Furthermore, we noticed several mitochondrial genes in the GTF were missing their chromosome ID, in turn preventing accurately calculating cell percentage mitochondrial content downstream; mitochondrial genes were therefore also prefixed in the GTF (MT-). Finally, we annotated the key neural crest specifier Snai2 in the GTF (ENSGALG00000030902). Reads were processed and aligned to GalGal6 using CellRanger 4.0.0 (10x Genomics).

The aligned reads from CellRanger (BAM) were re-counted using Velocyto (La Manno et al., 2018) to generate spliced and unspliced counts for each gene, which are used in subsequent RNA velocity analysis.

### Downstream pre-processing

All downstream analysis was wrapped into a custom Nextflow pipeline to simplify re-running the analysis by handling parallel processing of data subsets.

#### Data filtering

Quality control, data filtering, cell clustering, dimensionality reduction and initial data visualisation was carried out using Seurat v4.0.5 (Hao et al., 2021). Data filtering was carried out on each sequencing batch separately before data integration. To ensure the removal of poor-quality cells whilst also minimising over-filtering, we first applied a medium filtering threshold (cells expressing fewer than 1000 genes, more than 6500 genes, or had more than 15% mitochondrial content were excluded from subsequent analysis) followed by the removal of remaining poor-quality cell clusters.

#### Dimensionality reduction and clustering

Linear dimensionality reduction was carried out using principal component analysis (PCA). The number of principal components (PCs) used to construct the subsequent shared-nearest-neighbour (SNN) graph was determined unbiasedly, using an approach developed by the Harvard Chan Bioinformatics Core (HBC). First, we calculate the PC at which subsequent PCs contribute less than 5% of standard deviation, and the point where the cumulative variation of PCs exceeds 90%. The larger of these two values is selected. Second, we calculate the PC where the inclusion of consecutive PCs contributes less than 0.1% additional variation. The smaller value from these two steps is selected as a cut-off.

Louvain clustering was carried out with varying resolutions depending upon the data subset in question. A high clustering resolution (resolution=2) was used to identify poor quality clusters and contaminating cell states. This was to ensure that only poor-quality cells were filtered from the dataset. All other clustering was carried out using a resolution of 0.5.

#### Filtering of poor-quality clusters

Cell quality was measured using unique gene and UMI counts. Clusters in which both metrics fell below the 25th percentile relative to the rest of the dataset were considered poor quality and excluded from further analysis.

#### Data integration

After the removal of poor-quality cells, cells from the two separate batches were integrated into a single full dataset (Figure 2–figure supplement 1H-I). To avoid over-correction and removal of stage differences, we integrated the data from the two sequencing batches using STACAS v1.1.0 (Andreatta & Carmona, 2021). This R package allows for alignment of datasets containing partially overlapping cell states.

#### Regressing out confounding variables

Before downstream analysis, we regressed out percentage mitochondrial content, cell sex, and cell cycle. After regressing out percentage mitochondrial content, we identified a residual sex effect whereby cells clustered according to the expression of either W or Z chromosome genes. This resulted in a duplication of cell clusters, most apparent at HH4 (Figure 2–figure supplement 1A-D). Cells were k-means clustered according to their expression of W chromosome genes and classified as either male or female. This classification was used to regress out cell sex. Cell cycle was observed to have a strong effect on cell clustering and was therefore also regressed out during data scaling (Figure 2–figure supplement 1E).

#### Removal of contaminating clusters

We found small contaminating clusters of primordial germ cells, blood islands, mesoderm and endoderm within the dataset (Figure 2–figure supplement 1F, G). To semi-unbiasedly remove these clusters, we calculated the average expression of candidate markers for each of these cell states and removed clusters in which the median expression of any of these modules was greater than the 90^th^ percentile relative to the rest of the dataset (Figure 2–figure supplement 1F’’).

### Downstream analysis

#### Unbiased cell classification

Cell states were classified on each stage independently using a binary knowledge matrix (Supplementary File 1). First, the full dataset was split by stage and then re-scaled and re-clustered as described above. Our binary knowledge matrix consists of binarized expression of 76 genes across 24 cell states. For each gene, published *in situ* hybridisation expression patterns were used to determine expression within each defined cell state. Given that gene expression can vary dramatically between stages, and some cell states are known to not be present at specific timepoints (i.e., delaminating neural crest at HH4), we restricted the possible cell states at specific stages. At HH4 there were 7 possible cell states, including non-neural ectoderm (NNE), node, streak, extra-embryonic (EE), early NPB (eNPB), early neural (eN), early caudal neural (eCN). At HH5 there were 17 potential states, this included all cell states in the binary knowledge matrix, except for the later counterparts to the earlier cell states which are ventral forebrain (vFB), forebrain (FB), midbrain (MB), hindbrain (HB), neural crest (NC), delaminating neural crest (dNC) and presumptive epidermis (pEpi). At HH6 we included the same cell states as HH5 except for EE and eNPB given that no cell clusters were classified as such at HH5. For stages HH7, ss4 and ss8 we included 17 cell states, excluding the following states which are only found at earlier stages (NNE, node, streak, eN, eCN, eNPB, EE).

For each cell, we calculated the average scaled expression of the genes expressed in each cell state according to our binary knowledge matrix. Cell clusters were then assigned a cell state, depending on which cell state exhibited the highest median expression in that given cell cluster. Assigned cell states at each of the 6 sampled developmental stages were then transferred to the full dataset.

#### Gene module identification and filtering

To calculate groups of highly correlated genes, we first filtered genes which do not correlate (Spearman correlation < 0.3) with at least 3 other genes. Then we calculated gene modules using the Antler R package (Delile et al., 2019), which iteratively clusters genes hierarchically based on their gene-gene Spearman correlation. After each iteration, gene modules were filtered to remove gene modules which are not highly expressed (expressed in fewer than 10 cells) or which do not show consistent high expression across genes (fewer than 40% of genes in the module are expressed after binarization). Gene modules were calculated on different subsets of cells: the full dataset (Figure 4E and Figure 4–figure supplement 1A), ss8 (Figure 5A and Figure 5–figure supplement 1A), and the NPB subset (Figure 7E). Calculating gene modules on different subsets allowed us to zoom in on specific aspects of ectodermal lineage segregation.

Many genes correlate across the dataset, but do not show differential expression between cell states and therefore do not provide useful information regarding lineage segregation. Therefore, to identify which gene modules correlate with lineage segregation, modules were filtered to only include those in which more than 50% of their genes were differentially expressed (logFC > 0.5, adjusted p-value < 0.001) in at least one cell state relative to the rest of the dataset. For the NPB subset, this filtering of differential expression was just between the neural crest (NC, dNC) and placodal (aPPR, pPPR, PPR) states rather than between all cell states (logFC > 0.25, adjusted p-value < 0.001).

To remove gene modules which correlate with technical variation, we filtered those in which more than 50% their genes showed differential expression between sequencing batches (logFC > 0.25, adjusted p-value <0.001).

For the gene modules calculated on the full dataset, we were particularly interested in gene modules that segregated into one of the three lineages (neural, neural crest or placodal) at later stages. To focus our analysis on these gene modules, we applied an additional filtering step to only keep gene modules in which more than 50% of genes were differentially expressed between one of the three lineages at ss8 (logFC > 0.25, adjusted p-value <0.001). After filtering 9 gene modules remained (Figure 4–figure supplement 1A, Figure 4–Source Data 1). Modules were further manually filtered for visualisation (Figure 4E).

For the gene modules calculated on the NPB subset we wanted to focus on the co-expression and sequential activation of gene modules that contain well-characterised neural crest and PPR markers. We therefore further filtered NPB gene modules to include only those that had genes that were defined within neural crest, delaminating neural crest or PPR cell states in the binary knowledge matrix. After this final filtering step, 7 gene modules remained (Figure 7E, Figure 7–Source Data 1).

#### Identification of undetermined cells at the NPB

To identify which cells expressed competing pan-placodal and pan-neural crest modules at similar levels, we calculated the average expression (normalised counts) of these two gene modules in each cell and then ordered cells by their ratio of pan-placodal to pan-neural crest module expression. Cells were then grouped into 10 evenly sized bins and visualised on UMAPs.

#### Co-expression distribution plots

To investigate the distribution of co-expression between cells, co-expression histograms were plotted (Figure 5I). Co-expression values were calculated by multiplying the average normalised expression of the pan-placodal and pan-neural crest gene modules.

#### Co-expression visualisation

The average normalised expression of each gene module was scaled between 0 and 1. A 100×100 two-colour matrix was created using Seurat’s BlendMatrix function. Each cell was assigned a colour from this matrix based on their scaled average expression of the gene modules (Figure 5H). Only cells which express both modules above a threshold of 0.3 after scaling were visualised.

#### RNA velocity

Spliced and unspliced matrices obtained from Velocyto (La Manno et al., 2018) (see *Read Alignment*) were analysed using scVelo (Bergen et al., 2020) to generate cell velocity vectors. Velocyto matrices were intersected with each of the data subsets generated in Seurat. First and second order moments were then calculated for each cell across its 20 nearest neighbours using the top 20 PCs. RNA velocity was ran using the dynamical model whilst accounting for differential kinetics; this allows for differential splicing rates between genes as well as differential splicing kinetics between cell states.

Latent time was calculated and used downstream as a measure of a cell’s internal clock. When calculating latent time, cells were rooted to the earliest timepoint present in the data subset; in cases where only one stage was present, the cell root was calculated automatically. For all latent time estimates, we used a diffusion weighting of 0.2.

#### Modelling gene expression dynamics

Lineage inference was calculated using CellRank, which uses a probabilistic approach to assign cell lineage (Lange et al., 2022). Cell transition matrices were calculated using RNA velocity (0.8 weighting) and nearest neighbour graph (0.2 weighting) kernels. A softmax scale of 4 was set to prevent transitions tending towards orthogonality. When predicting cell transitions, terminal states were specified according to the lineages which we were wanting to model gene dynamics across. For the full dataset, we predicted cell transitions towards either neural, placodal or neural crest lineages. Terminal states were specified as: delaminating neural crest and neural crest cell states at ss4 and ss8 for the neural crest lineage; hindbrain, midbrain, and forebrain cell states at ss4 and ss8 for the neural lineage; anterior PPR, posterior PPR and PPR cell states at ss4 and ss8 for the placodal lineage (Figure 4–figure supplement 1C). For our NPB subset we predicted cell transitions towards either placodal or neural crest lineages, with terminal states specified as: delaminating neural crest and neural crest cell states at ss8 for the neural crest lineage; aPPR and PPR cell states at ss8 for the placodal lineage.

Gene expression dynamics were calculated by fitting general additive models (GAM), which modelled scaled gene expression as a function of latent time using the mgcv package in R. Lineage absorption probability obtained from CellRank were used to weight a GAM for each of the predicted lineages. To generate a smoothed prediction of gene expression change, a cubic spline was fitted with knots=4. When modelling the dynamics of entire gene modules, all genes within a given module were included in the GAM.

#### Modelling mediolateral gene expression patterns from scRNAseq data

<u>Following principal component analysis of cells at HH7, the order of cell states across the inverse of PC 1 were found to reflect the M-L positioning of cells *in vivo*. Genes which were. To model the expression of placodal and neural crest markers across the M-L axis, GAMs were fitted for the scaled expression of each gene across PC 1 using ggplot2::geom_smooth</u>.

### Fluorescent *in situ* hybridisation chain reaction

Hybridisation chain reaction v3 (HCR) was carried out as previously described (Buzzi et al., 2022; Choi et al., 2018). Embryos were fixed in 4% PFA for 1h at room temperature, dehydrated in a series of methanol in PBT, and stored overnight at −20°C. Samples were rehydrated and treated with proteinase-K for 3 min (20 mg/mL). Next samples were post-fixed in 4% PFA for 20 min and sequentially washed on ice for 5 min in PBS, 1:1 PBT/5×SSC (5×sodium chloride sodium citrate, 0.1% Tween-20), and 5×SSC. Samples were then pre-hybridised in hybridisation buffer for 5 min on ice, followed by 30 min at 37°C. Next, samples were hybridised overnight at 37°C with HCR probes in hybridisation buffer (4 pmol/mL). Excess probe was washed off with 15 min washes in probe wash buffer at 37°C, before preamplification in amplification buffer at room temperature for 5 min. Hairpins (30 pmol) were incubated at 95°C for 90 s and left to cool to room temperature for 30 min before being added to the amplification buffer. Samples were then incubated with the hairpin/amplification buffer solution overnight at room temperature. Excess hairpins were washed off with two 5 min and two 30 min washes in 5×SSC. After a 5 min incubation in DAPI (10 mg/mL), samples were washed 3 times for 10 min in 5×SSC before being imaged using a Leica SP5 laser scanning confocal inverted microscope using the LAS AF software.

Intensity measurements were calculated by taking the sum of slices from the image Z-stacks taken at three anteroposterior levels in ImageJ. The three intensity measurements were then independently scaled (Z-score) for each gene to allow relative measurements of gene expression across the anteroposterior axis. Transverse virtual sections across the anteroposterior axis were obtained using the Imaris 9.3.1 software. 10 μm sections were generated with the orthogonal slice tool.

## Data availability and reproducibility

10x single cell RNAseq was carried out in two batches and is available under two separate accession numbers (ArrayExpress: E-MTAB-10408 and E-MTAB-1144). Our NGS alignments and downstream analysis have been wrapped into custom Nextflow pipelines allowing for full reproducibility. For the code used in this analysis, including links to our Docker containers, see our GitHub repository at https://github.com/alexthiery/10x_neural_plate_border.

## Acknowledgements

We thank Teresa Rayon for assistance with establishing the single cell dissociation protocol and the Advanced Sequencing Facility at the Francis Crick Institute for the single cell RNA sequencing. We would also like to thank Igor Adameyko and Artem Artemov for guidance on co-expression analysis, Chantal Hubens for technical support, Alessandra Vigilante and Sami Leino for comments on the manuscript, and Streit and Luscombe lab groups for helpful discussions. This work was supported by the Wellcome Trust (108874/B/15/Z), the BBSRC (BB/S005536/1; BB/R006342/1) and in part by the Francis Crick Institute which receives its core funding from Cancer Research UK (FC001051), the UK Medical Research Council (FC001051), and the Wellcome Trust (FC001051). For the purpose of Open Access, the author has applied a CC BY public copyright license to any Author Accepted Manuscript version arising from this submission.

## Author contributions

A.S., J.B., A.T. and A.L.B. designed study. A.T., A.L.B., E.H. and A.S. collected samples and processed for scRNAseq. A.T., C.C. developed analysis pipeline and carried out data alignment. A.T. and E.H. carried out the downstream computational analysis of scRNAseq data, while A.L.B. and A.T. performed HCR analysis. A.T., A.L.B., E.H., N.L., J.B. and A.S. interpreted the results. A.T., A.L.B., E.H. and A.S. made figures and wrote the manuscript. All authors provided feedback on the manuscript.

