## Supplementary Figures for "A gradient border model for cell fate decisions at the neural plate border"

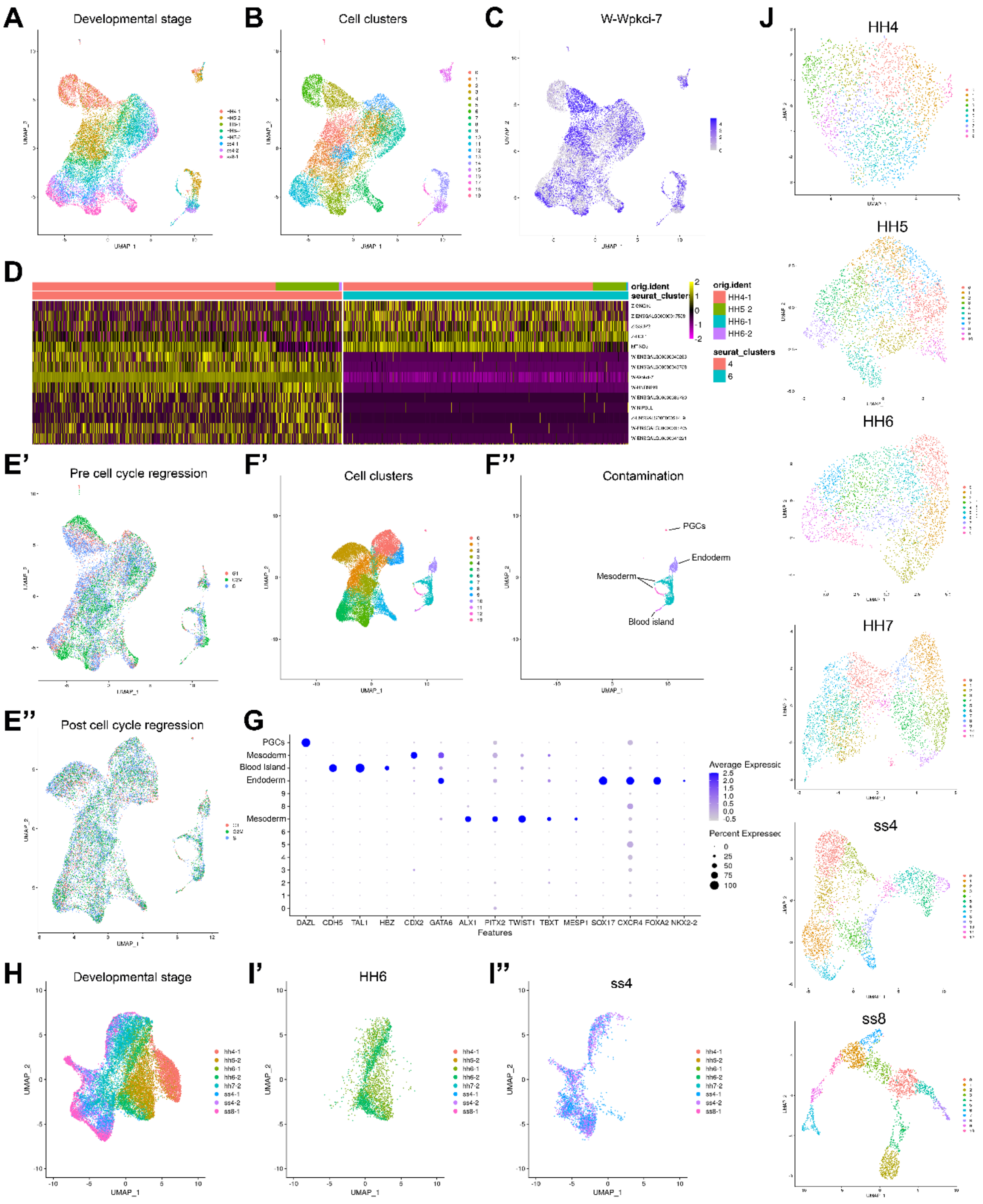


**Figure 2-figure supplement 1. (A-B)** UMAP of full dataset prior to regressing out confounding factors. Cells are coloured by developmental stage (A) and Seurat cell clusters (B). **(C)** Feature plot displaying W (male) sex-chromosome gene Wpkci-7, illustrating the cell sex effect after clustering. **(D)** Heatmap of selected sex genes across clusters 4 and 6. These clusters contain mostly cells from HH4 and highlight the clear sex effect of W and Z chromosome genes between these clusters. **(E)** UMAP of cells prior to (E') and post (E'') regressing out the cell cycle effect. **(F')** UMAP displaying cell clusters after regressing out sex and cell cycle effects. **(F'')** UMAP highlighting contaminating cell states following cell state classification. These include primordial germ cells (PGCs), mesoderm, endoderm, and blood islands. **(G)**

Dotplot for the expression of genes used for identification of contaminating cell clusters. **(H)** UMAP of all cells after filtering of contaminating populations and regressing out confounding variables. Cells are coloured by developmental stage. **(I)** UMAP plots showing successful integration of two sequencing batches at stages HH6 (I') and ss4 (I''). **(J)** UMAPs displaying cell clusters calculated at each developmental timepoint. Individual stages are subset from the full filtered dataset displayed in (H).

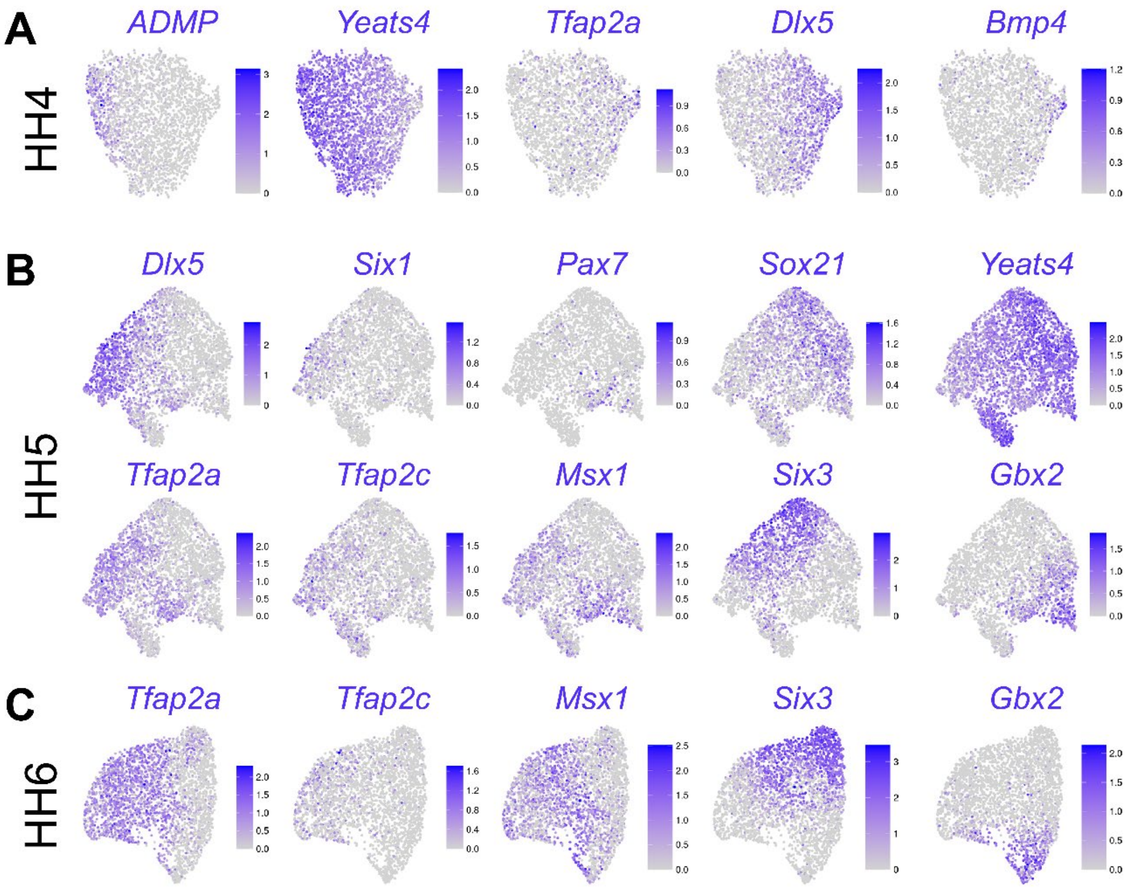


**Figure 2-figure supplement 2.** Feature plots showing expression of marker genes on UMAPs of developmental stages HH4 **(A)**, HH5 **(B)** and HH6 **(C).**

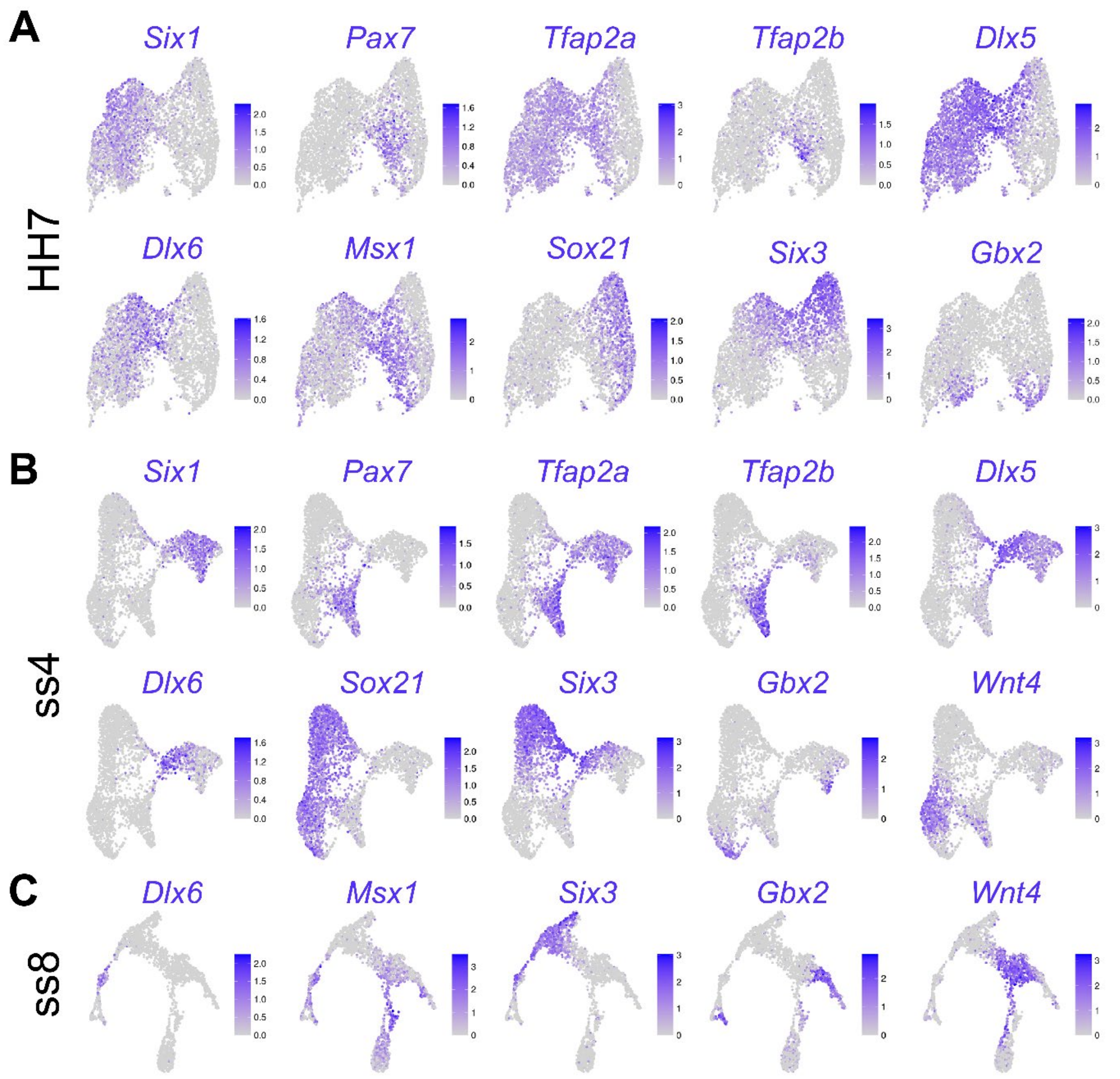


**Figure 3-figure supplement 1.** Feature plots showing expression of marker genes on UMAPs of developmental stages HH7 **(A)**, ss4 **(B)** and ss8 **(C).**

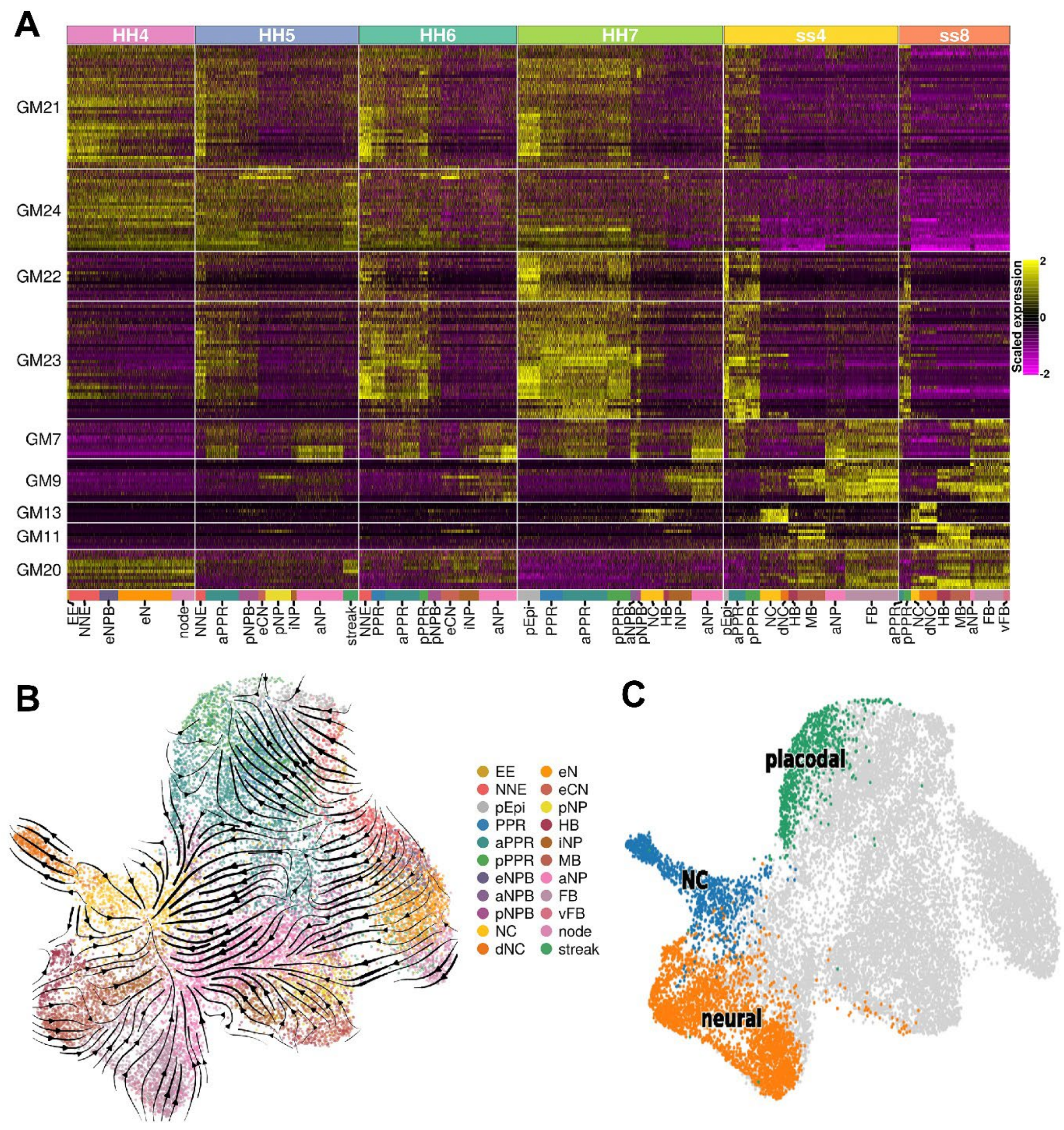


**Figure 4-figure supplement 1. (A)** Gene modules calculated across the full dataset (full list of gene modules available in Figure 4-Source Data 1). Gene modules were unbiasedly filtered to include those that only display differential expression between cell states and not between sequencing batches and then further filtered to include only those that are differentially expressed between the neural, neural crest and placodal cell states at ss8. 9 gene modules remained after filtering. **(B)** UMAP plot of the full dataset overlaid with RNA velocity vectors depicting the predicted directionality of transcriptional change. Cells coloured by cell state. **(C)** Selected terminal states (placodal, neural crest and neural) used for CellRank lineage inference.

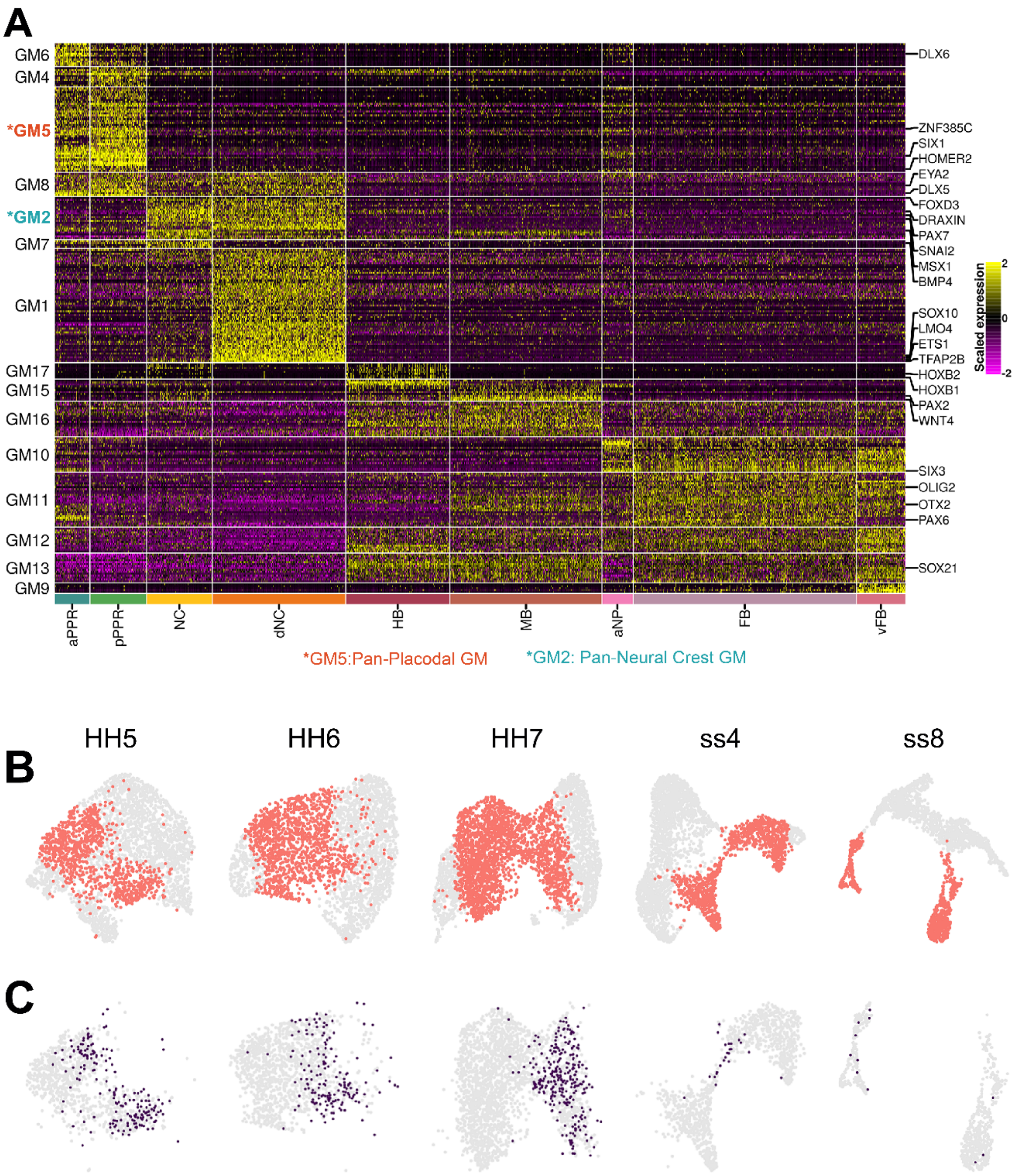


**Figure 6-figure supplement 1. (A)** Heatmap showing gene modules calculated from ss8 cells and then filtered to include only those that are differentially expressed between cell states (see methods; full list of gene modules available in Figure 5-Source Data 1). GM5 and GM2 are highlighted as the selected pan-placodal and pan-neural crest gene modules used for subsequent analysis in Figure 5. **(B)** UMAP plots of cells from HH5-ss8, with PPR, neural crest and NPB cells highlighted in red. These cells were selected as the ‘NPB subset’ in Figures 5 and 6. **(C)** Bin3 cells from Figure 5E mapped back to each individual stage, highlighting cells in the NPB subset with similar levels of co-expression of placodal GM5 and neural crest GM2.

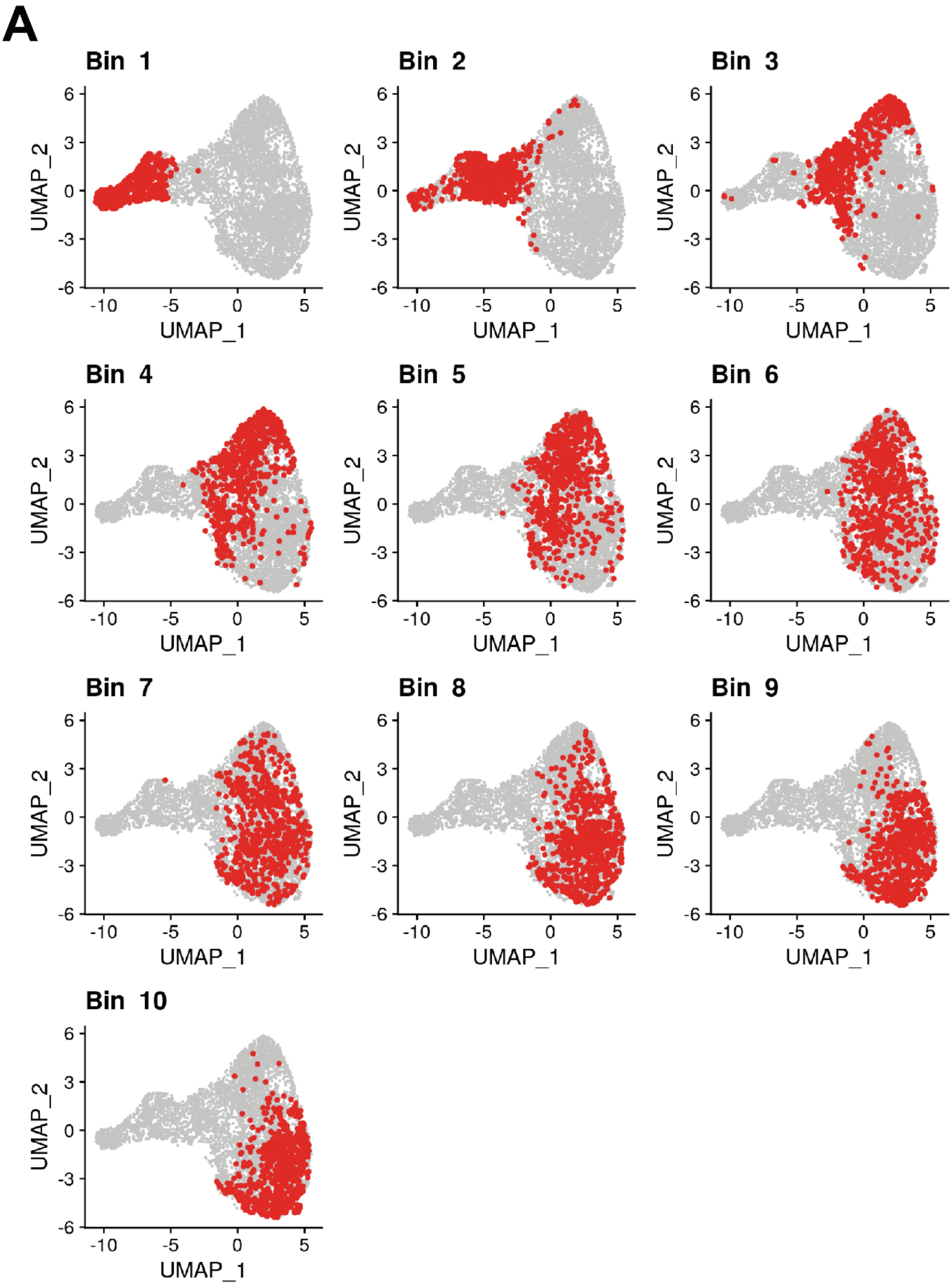


**Figure 6-figure supplement 2. (A)** UMAPs of the NPB subset. Cells from the NPB subset were ordered by their expression ratio of placodal and neural crest gene modules and then separated into 10 evenly sized bins (Figure 6D). Cells belonging to each of these 10 bins are displayed on the UMAPs.

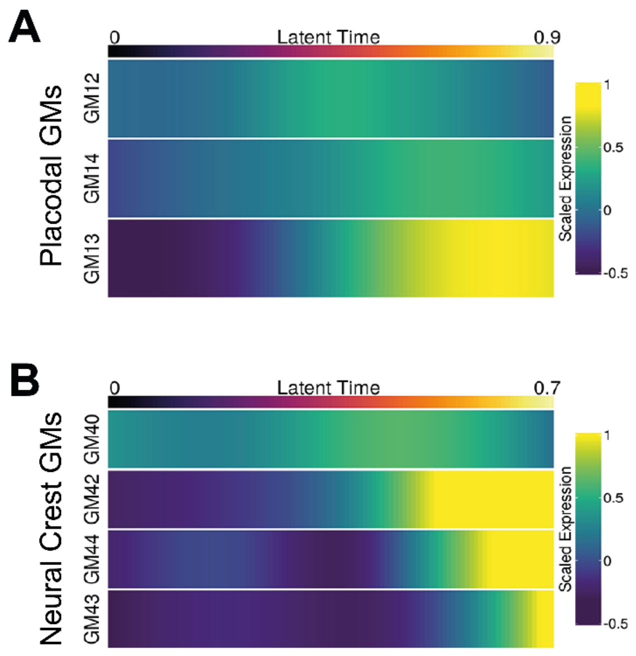


**Figure 7-figure supplement 1. (A)** Heatmap displaying predicted scaled expression of placodal gene modules (GM12, GM13 and GM14) within the placodal lineage across latent time. Gene expression was predicted by fitting a GAM which models scaled gene expression as a function of latent time. Gene expression was weighted in the GAM using the placodal fate lineage absorption probability. **(B)** Heatmap displaying predicted scaled expression of neural crest gene modules (GM40, GM42, GM43 and GM44) within the neural crest lineage across latent time. Gene expression was modelled as in (A) but using neural crest fate absorption probabilities to weight gene expression.
